# Interference Requirements of Type III CRISPR-Cas Systems from *Thermus thermophilus*

**DOI:** 10.1101/2023.09.03.556089

**Authors:** Karyna Karneyeva, Matvey Kolesnik, Alexei Livenskyi, Viktor Zgoda, Vasiliy Zubarev, Anna Trofimova, Daria Artamonova, Yaroslav Ispolatov, Konstantin Severinov

**Author notes:** Corresponding author. Waksman Institute, Rutgers, The State University of New Jersey, NJ 08854 USA.

## Abstract

Among the diverse prokaryotic adaptive immunity mechanisms, the Type III CRISPR-Cas systems are the most complex. The multisubunit Type III effectors recognize RNA targets complementary to CRISPR RNAs (crRNAs). Target recognition causes synthesis of cyclic oligoadenylates that activate downstream auxiliary effectors, which affect cell physiology in complex and poorly understood ways. Here, we studied the ability of III-A and III-B CRISPR-Cas subtypes from *Thermus thermophilus* to interfere with plasmid transformation. We find that for both systems, requirements for crRNA-target complementarity sufficient for interference depend on the target transcript abundance, with more abundant targets requiring shorter complementarity segments. This result and thermodynamic calculations indicate that Type III effectors bind their targets in a simple bimolecular reaction with more extensive crRNA-target base pairing compensating for lower target abundance. Since the targeted RNA used in our work is non-essential for either the host or the plasmid, the results also establish that a certain number of target-bound effector complexes must be present in the cell to interfere with plasmid establishment. For the more active III-A system, we determine the minimal length of RNA-duplex sufficient for interference and show that the position of this minimal duplex can vary within the effector. Finally, we show that the III-A immunity is dependent on the HD nuclease domain of the Cas10 subunit. Since this domain is absent from the III-B system the result implies that the *T. thermophilus* III-B system must elicit a more efficient cyclic oligoadenylate dependent response to provide the immunity.

## Introduction

Prokaryotic CRISPR-Cas defense systems comprise CRISPR arrays consisting of two or more identical repeats separated by unique spacers and associated clusters of *cas* genes [1,2]. CRISPR-Cas systems are encoded by ∼85% of archaeal and ∼40% bacterial genomes [1] and multiple examples of CRISPR-Cas systems encoded by viruses that infect prokaryotes have been reported [3]. The CRISPR-Cas defense can be divided into three stages. At the adaptation stage, short DNA sequences originating from mobile genetic elements such as viruses or plasmids are inserted into CRISPR arrays forming new spacers. Spacer insertion is accompanied by a duplication of the repeat sequence [4,5]. At the expression stage, CRISPR arrays are transcribed, and the transcripts are processed into small CRISPR RNAs (crRNAs) that contain repeat-derived and spacer-derived segments [6–11]. At the interference stage, individual crRNAs interact with Cas proteins forming effector complexes that recognize protospacers, sequences complementary to crRNAs spacer segments. Protospacer recognition by the effector complex either directly leads to destruction of the genetic invader’s nucleic acid or leads to changes in cellular physiology that limit the spread of the invader in the host population.

Based on the structures of *cas* gene clusters, two classes of CRISPR-Cas systems that comprise six types and numerous subtypes have been recognized [1]. Type I and Type IV effectors of Class 1 systems and Type II and Type V effectors of Class 2 systems recognize DNA. Since recognition of spacers in CRISPR arrays must be avoided to prevent auto-immunity, effectors of these DNA-recognizing systems rely on an additional element, protospacer adjacent motif (PAM) to distinguish self (a spacer in CRISPR array) from non-self (a protospacer in foreign DNA) sequences. PAMs are specifically recognized, in a double-stranded form, by one of the subunits of multisubunit Class 1 effectors or a dedicated domain of a single-subunit Class 2 effectors. PAM recognition licenses partial melting of target protospacer DNA and its interrogation by the crRNA spacer segment for complementarity [12–14]. Initial base pairing between crRNA and the target in its PAM-proximal part is critical for the formation of a ternary complex needed to initiate interference/target destruction. Even single mismatches in this part, referred to as “seed”, strongly diminish or abolish the ternary complex formation and, therefore, interference [15]. Although precise locations of “seed” regions may be context-dependent, approximate positions, depending on the system, extend for about 6-12 base pairs [10,16–18]. For example, in the case of a well-studied *E. coli* Type I-E Cascade effector, the seed region extends for eight PAM-proximal nucleotides. However, the sixth position is insensitive to mismatches as the crRNA base at this position is flipped out, interacts with a subunit of the effector and is thus unavailable for base pairing with the target protospacer [19,20]. Upon formation of a duplex in the seed, the RNA-DNA heteroduplex is extended over the entire length of the spacer segment of crRNA (20-32 bases depending on the system) [21]. Requirements for spacer-target complementarity outside the seed are less stringent and multiple mismatches can be tolerated with no apparent effects on interference [10,16–18].

In DNA-recognizing systems, the absence of PAM in CRISPR repeat sequences flanking spacers prevents auto-immunity. Type III (Class 1) and Type VI (Class 2) effectors recognize RNA molecules complementary to crRNA spacer segments, which solves the problem of auto-immunity. As a consequence, neither Type III nor Type VI systems require a PAM for target recognition. Yet, efficient interference by these systems requires a lack of complementarity between the target RNA and repeat-derived segments of crRNA (crRNA handle) [22–24]. Since the targets of these systems are single-stranded, there also does not appear to be a requirement for a seed, since base pairing between crRNA spacer and RNA target can in principle begin in any region (though likely should be affected by the secondary structure of the target). While Type VI systems are the least common, Type III systems are widespread and constitute, correspondingly, 34 and 25% of archaeal and bacterial CRISPR-cas loci [25]. Type III systems are also most complex mechanistically and many details of their function remain obscure [26]. Despite the fact that Type III effectors recognize RNA targets, these systems efficiently prevent infection with DNA viruses or transformation with plasmids from which RNAs complementary to crRNA spacers are transcribed. It is still not clear whether they directly destroy foreign DNA after its transcript is recognized or whether target RNA recognition causes cessation of cell growth indirectly, providing enough time to clear the foreign genetic element. RNA target recognition by widespread Type III-A and III-B effectors leads to production of cyclic oligoadenylates (cOAs) [27–29]. *In vitro*, cOAs stimulate auxiliary effectors, including nucleases containing cyclic oligonucleotide sensing domains [30]. It is thought that *in vivo* cOAs also act as secondary messengers and ensure that RNA target recognition by Type III effectors allows interference with foreign DNA [31–33]. For example, stimulation of non-specific cOA-activated RNases has global effects that prevent the development of DNA-containing foreign genetic elements and/or leads to infected cell death [34]. All Type III-A and many Type III-B effectors contain a subunit with an HD nuclease domain [1], which may directly attack DNA upon the recognition and binding of target RNA [23,35]. Eventually, the RNA protospacer recognized by a Type III effector is cleaved due to an RNase activity of subunits that interacts with the crRNA spacer segment [23,35]. This shuts off the cOAs synthesis and the nuclease activity of the HD domain when it is present [36] and, therefore, turns off the interference process.

Given the recent progress in the development of tools based on Type III effectors [37–41], it is important to understand their requirements for target recognition. Maniv et al. scanned the 35 nucleotide long crRNA spacer of the Type III-A *Staphylococcus epidermidis* system with mutations introducing 8-nucleotide long mismatches with a target transcribed from a conjugative plasmid and monitored the conjugation efficiency [42]. They found that only a mismatch in the most distal part of the duplex (positions +27 to +35, where +1 denotes the first, 5’-end proximal position of crRNA spacer) did not affect interference. In fact, even a small mismatch at positions +2 to +4 prevented the interference. They also tested a different crRNA-protospacer pair for ability to provide protection from phage infection. In this setup, none of the 8-nucleotide mismatches had any effect on the ability of cells to interfere with phage plaque formation. Larger mismatches involving either half of the spacer-protospacer heteroduplex abolished phage protection, however, leading to the appearance of plaques. Since plaque formation requires multiple cycles of reinfection and lowered phage burst sizes can lead to the absence of plaques, the authors also monitored the efficiency of infective center formation, an assay that determines whether an infected cell produces at least one infectious progeny phage particle. In this assay only a mismatch between positions +1 to +8 allowed productive infection, suggesting that the affected region may play a role of a seed. However, a smaller mismatch between positions +2 to +4 had no effect on infective center formation. The authors speculated that the apparent discrepancy between plasmid conjugation and phage infection assays results may be due to differences in the sequences of spacer-protospacer pairs, differences in locations of targeted protospacers in mobile genetic elements or levels of their transcription, or differences in the mechanisms of entry of plasmid and phage DNA in immune cells.

Pyenson et al. studied target recognition requirements of the *S. epidermidis* Type III-A system by introducing random mutations into 10-nucleotide segments of plasmid-borne protospacer [43]. No single mismatches abolishing immunity were detected. However, interference with some plasmids harboring four mismatches within the first 10-nucleotide segment of crRNA was decreased. Six or more mismatches in the second segment were required to affect interference, while plasmids containing ten mismatches in the last segment (and thus lacking any complementarity with the 3’-end proximal part of crRNA) were subject to interference.

Staals et al. studied requirements for target RNA cleavage by *Thermus thermophilus* Type III-A effector *in vitro* [44] by introducing single-nucleotide mismatches in protospacer region corresponding to the first seven spacer nucleotides proximal to the 5’ repeat-derived crRNA handle. They also scanned the crRNA-target duplex with 5-nucleotide mismatches. Target cleavage is catalyzed by the Csm3 subunits of the effector. With a fully matching target, cleavages between positions +5/+6, +11/+12, +17/+18, +23/+24, +29/+30, and +35/+36 at the boundaries of segments recognized by individual Csm3 subunits were observed. The mismatches hampered target RNA cleavage at positions located immediately downstream but had no effect on other cleavage sites, indicating that all mismatched targets were bound by the effector. The authors therefore suggested that the Type III-A *T. thermophilus* effector lacks a seed region.

Target recognition requirements of *T. thermophilus* Type III-B effector were studied by Steens et al. [37]. The large Cmr2 subunit of this effector lacks the HD domain indicating that in *T. thermophilus*, the III-B immunity relies only on target RNA recognition, activation of auxiliary effectors by cOAs production (or some other, yet unknown mechanism), followed by target RNA cleavage. The authors used an *in vitro* assay measuring the rate of pyrophosphate production during the conversion of ATP to cOAs upon target recognition to localize target regions crucial for cOAs synthesis. They found that single mismatches between the target and crRNA in a region corresponding to the first seven spacer nucleotides had no effect on target cleavage and, therefore, did not interfere with target recognition. However, mismatches at positions +1 and +2 greatly diminished the production of cOAs. The affected region was designated as Cas10-activating region (CAR). They also studied the effects of 5-nucleotide mismatches in the crRNA-target duplex. Only mismatches at positions +19/+23 and +25/+29 affected target recognition as judged by EMSA.

The effects of crRNA-target mismatches on the cOAs synthesis were also studied *in vitro* by Nasef et al. using a *S. epidermidis* Type III-A effector preparation loaded with three different crRNAs [45]. They used an effector containing a mutant Csm3 subunit that was unable to perform target cleavage, thus ensuring continuous production of cOAs upon target binding. They found that a triple mismatch between positions +1/+3 abrogates cOAs synthesis. Since target binding was not affected, the mismatched region therefore corresponded to CAR. Single mismatches at positions +2 and +7 also abolished cOAs production upon target binding. Curiously, the corresponding mismatches had no effect on cOAs synthesis when another spacer-protospacer pair was tested. *In vivo,* the +1/+3 mismatches in either spacer-protospacer pair had no effect on interference with plasmid transformation.

Studies of target recognition requirements of Type III effector complexes are further complicated by the fact that maturation results in crRNAs with constant 5’ and variable 3’ ends [46,47]. The lengths of mature crRNAs vary with 6 nucleotide periodicity [44,48–50]. The Csm3/Cmr4 subunits interact with the 6-nucleotide segments of the crRNA spacer and effector variants with different numbers of Csm3 (for III-A systems) and Cmr4 (for III-B systems) subunits have been reported [51,52]. Thus, the lengths of Csm3/Cmr4 filaments serve as rulers defining the length of bound crRNAs. To exclude the effects of effector complex heterogeneity, Steens et al. [37] reconstituted effectors with crRNA molecules of defined lengths and tested them for recognition of mismatched targets. In experiments with the *T. thermophilus* III-B effector charged with full-length 46-nucleotide crRNA, the cOAs production was inhibited by 5-nucleotide mismatches at Cmr4-bound segments +1/+5, +7/+11, and +19/+23. The +19/+23 mismatch also abolished cleavage of target RNA suggesting that a putative “seed” region overlaps with the segment recognized by the fourth Cmr4 subunit in the filament.

In the III-B effector lacking one Cmr4 unit and charged with shorter, 40-nucleotide crRNA [52], the cOAs production was inhibited by mismatches between duplex segments +1/+5, +7/+11, +13 /+17, and +19/+23. Target cleavage was affected by +13 /+17 and +19/+23 mismatches, indicating that with shorter crRNA both the third and the fourth Cmr4 subunits became essential for target recognition, effectively leading to expansion of the seed.

Here, we studied target recognition requirements by Type III-A and III-B CRISPR-Cas systems from *T. thermophilus* HB27. Although *T. thermophilus* III-A and III-B effectors are composed of different proteins [25], they rely on a common set of CRISPR arrays as a source of crRNAs [53]. The Type III immunity of *T. thermophilus* HB27 is highly active and strongly decreases plaque formation by DNA viruses when crRNAs are complementary to viral transcripts. Both the III-A and III-B systems alone protect cells from viral infection [53]. At least the III-A system, alone, inhibits transformation with plasmids containing sequences complementary to crRNA spacers and the level of protection correlates with the amount of transcripts produced from the targeted plasmid sequence [54]. The genome of *T. thermophilus* HB27 encodes three proteins containing cOA-sensing CARF domains: Csx1, Csm6, and Can1. *In vitro,* the first two proteins behave as cOA-activated RNases, while Can1 relaxes supercoiled plasmids and may thus be a cOA-dependent nickase [31,38,55]. The role of CARF domain nucleases, if any, in Type III immunity of *T. thermophilus* is unknown. In this work, we systematically defined target recognition requirements of both systems and determined the minimal length of continuous RNA duplex necessary for target recognition and interference by the III-A system. We also accessed the roles of CARF RNases and of the Type III-A largest effector subunit HD domain in the ability of immune cells to withstand plasmid transformation.

## Results

### Determining the boundaries of crRNA-target duplex sufficient for III-A and III-B interference

To study base pairing requirements of crRNA spacer for *T. thermophilus* III-A and III-B interference, the previously described strains lacking the *cmr4* gene coding for the III-B effector subunit or the *csm3* gene coding for the III-A effector subunit were used [53]. Below, we refer to these strains as III-A^+^ and III-B^+^, correspondingly. To monitor Type III immunity in either strain, two derivatives of the pMK18 vector plasmid [56] containing a protospacer (PS) corresponding to the 1^st^ spacer of the 8^th^ *T. thermophilus* HB27 CRISPR array were used [54]. The two plasmids differed in the orientation of the protospacer cloned into the pMK18 polylinker site downstream of a strong constitutive PslpA promoter. In the PS_dir plasmid, the PslpA-initiated transcripts are complementary to the spacer part of the crRNA. In the PS_rev plasmid the protospacer part of the PslpA-initiated transcript has the same sequence as the crRNA spacer. Based on high throughput RNA sequencing data, the abundance or reads corresponding to pMK18 transcripts initiated from PslpA exceeded the abundance of antisense transcripts by ∼80-fold in wild-type, III-A^+^, or III-B^+^ *T. thermophilus* strains (Supplementary Fig. S1). The antisense transcription is presumably due to activity of an unidentified divergent promoter.

The pMK18 control and protospacer-containing plasmids were transformed in III-A^+^ and III-B^+^ strains and aliquots of transformation mixtures and their serial dilutions were spotted on the surface of kanamycin-containing medium to select for transformants. In agreement with earlier data [54], both PS_dir and PS_rev were poorly transformed in the III-A^+^ strain and were thus subject to III-A interference (Fig. 1a). Both plasmids were also subject to III-B interference (Fig. 1b). For both PS_dir and PS_rev in the case of the III-A^+^ strain and for the PS_dir plasmid in the case of the III-B^+^ strain, rare kanamycin-resistant colonies were observed when undiluted transformation mixtures were spotted on selective medium. Genomic sequencing of randomly chosen kanamycin-resistant clones revealed point mutations in essential *cas* genes (Supplementary Table S2) or deletions of entire III-A (for the III-A^+^ strain) or III-B (for the III-B^+^ strain) *cas* operons (Supplementary Fig. S2 and Supplementary Table S2).

**Figure 1.**
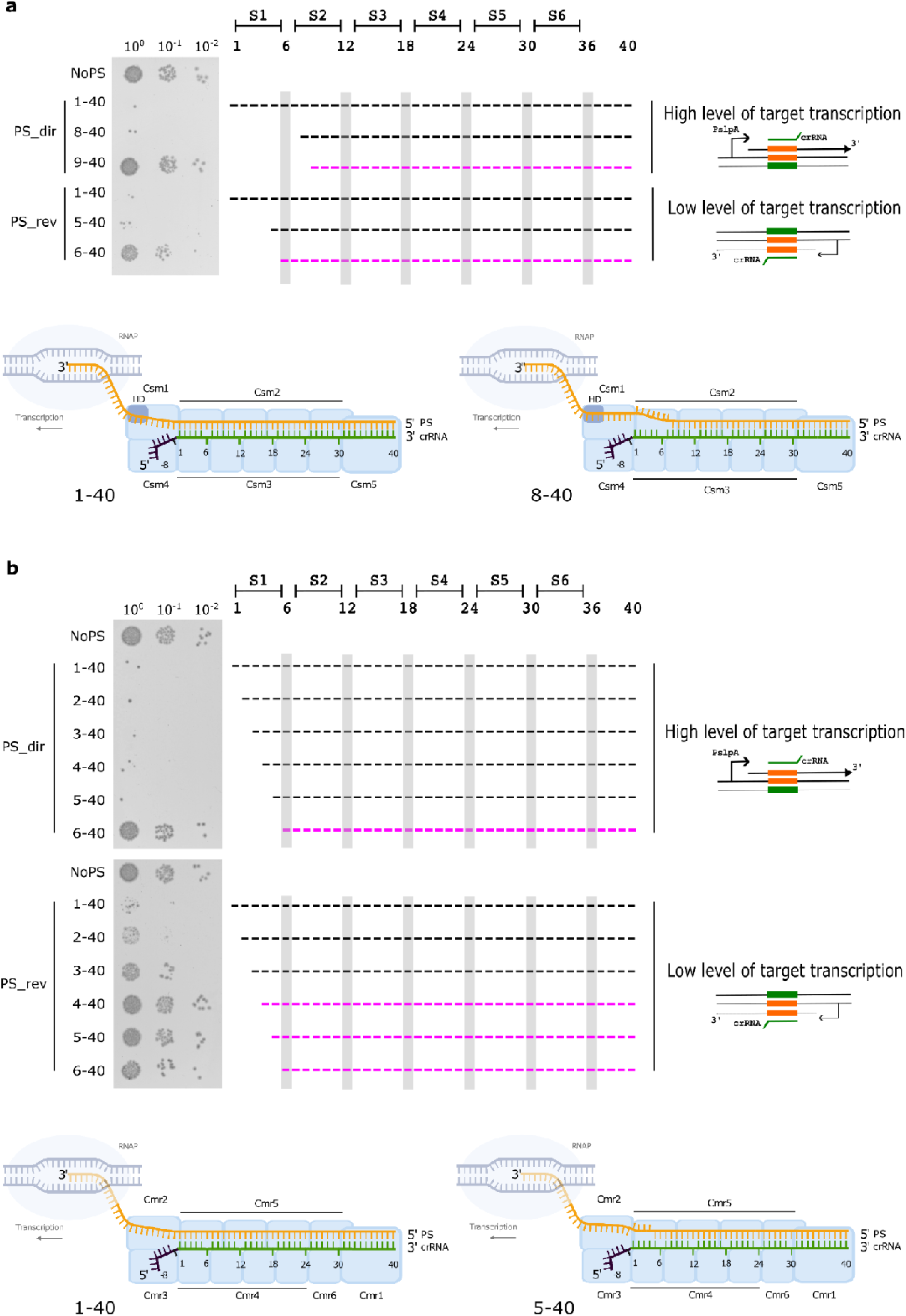
Defining spacer-protospacer complementarity requirements at the 5’ end-proximal part of the crRNA spacer for *T. thermophilus* Type III-A (a) or III-B (b) interference with plasmid transformation. *T. thermophilus* strains harboring active III-A (**a**) or III-B (**b**) CRISPR-Cas systems were transformed with the pMK18 vector plasmid (“NoPS”), or derivatives carrying a protospacer matching a Type III spacer. In cells carrying the PS_dir plasmid, the abundance of protospacer transcripts complementary to host crRNA and initiated from a strong PslpA promoter is ∼80 higher than in cells carrying PS_rev, where transcription is initiated from unidentified oppositely oriented promoters as schematically shown on the right. 10-μl aliquots of serial 10-fold transformation mixtures dilutions were dropped on kanamycin-containing agar plates where only cells harboring pMK18-based plasmids form colonies. Representative images of overnight growth at 65°C obtained in at least three independent experiments are shown. Results of transformation with PS_dir and PS_rev derivatives introducing spacer-protospacer mismatches are also shown. These derivatives are labeled to indicate the maximal expected lengths of the RNA duplex in effector-target complexes. PS_dir and PS_rev derivatives that are interfered with are indicated in black, those that are not subject to interference - in magenta. In schemes shown in the middle, the positions of the spacer (from +1 to +40) are labeled at the top, with “S1” through “S6” denoting crRNA segments that interact with individual Csm3 (III-A) or Cmr4 (III-B) effector subunits. Positions corresponding to crRNA bases +6, +12, +18, +24, and +30 are highlighted by gray vertical stripes. These bases are flipped and do not interact with the target (Supplementary Figure S5). At the bottom of each panel, schematic structures of effector complexes with a fully matching (left) and with maximally mismatched highly transcribed target (right) that support interference are shown.

We created versions of PS_dir and PS_rev plasmids with mutations introducing mismatches between the 5’ end-proximal portion of crRNA spacer and protospacer-containing transcripts, thus shortening the length of the crRNA-target duplex, and tested them for efficiency of transformation. A mismatch over the first 7 nucleotides of the crRNA-target duplex (an RNA duplex extending over positions +8/+40) had no effect on III-A interference against PS_dir (Fig. 1a). Shortening the duplex by an additional base pair, to +9/+40, abolished interference. For PS_rev, a +5/+40 but not the shorter +6/+40 duplex was sufficient for interference. In the case of the III-B^+^ strain, interference against the PS_dir plasmid disappeared when the duplex boundary was moved from position +5 to +6 (Fig. 1b). Thus, with respect to the upstream RNA duplex boundary, the interference requirements of the III-A system against the poorly transcribed PS_rev target are the same as those of the III-B system against the highly transcribed PS_dir target. In the case of PS_rev, a 3-residue mismatch prevented interference by the III-B system, a two-nucleotide mismatch decreased transformation efficiency ∼10-fold compared to the pMK18 control, while a plasmid with a single mismatch at position +1 transformed just slightly better than parental PS_rev as judged by the increased number of small colonies formed with undiluted transformation mixtures aliquots (Fig. 1b).

Since importance of the 3’ end-proximal part of crRNA spacer was studied only for the *T. thermophilus* III-A system [54], we repeated transformation experiments with PS_dir and PS_rev plasmids harboring nested multiple mismatches in the corresponding part of the protospacer with the III-B^+^ strain (Fig. 2). For PS_dir, mismatches up to position +31 (a +1/+30 duplex with the target) had no effect on interference (compared to +1/+23 in the case of the III-A system [54]). Mismatches extending up to positions +30 and +29 had a partial effect with substantial amounts of small colonies formed when the undiluted transformation mixtures were spotted on the selective medium. Duplexes of 27 base pairs or less were not sufficient for interference. In the case of PS_rev, the requirements were more strict and a +1/+37 or longer duplex (compared to +1/+26 in the case of the III-A system, [54]) was needed for interference. We note that the III-B effector purified from *T. thermophilus* cells is a 2.5:1 mixture of species containing crRNAs with 32 and 38 nucleotides spacer segments [52]. Presumably, introduction of mismatches beyond position +36 affects only effectors containing the longer crRNA. For *T. thermophilus* III-A effector purified from cells, a broad and rather indistinct distribution of crRNAs lengths was determined with the most abundant species corresponding to 39-nucleotide spacer [44].

**Figure 2.**
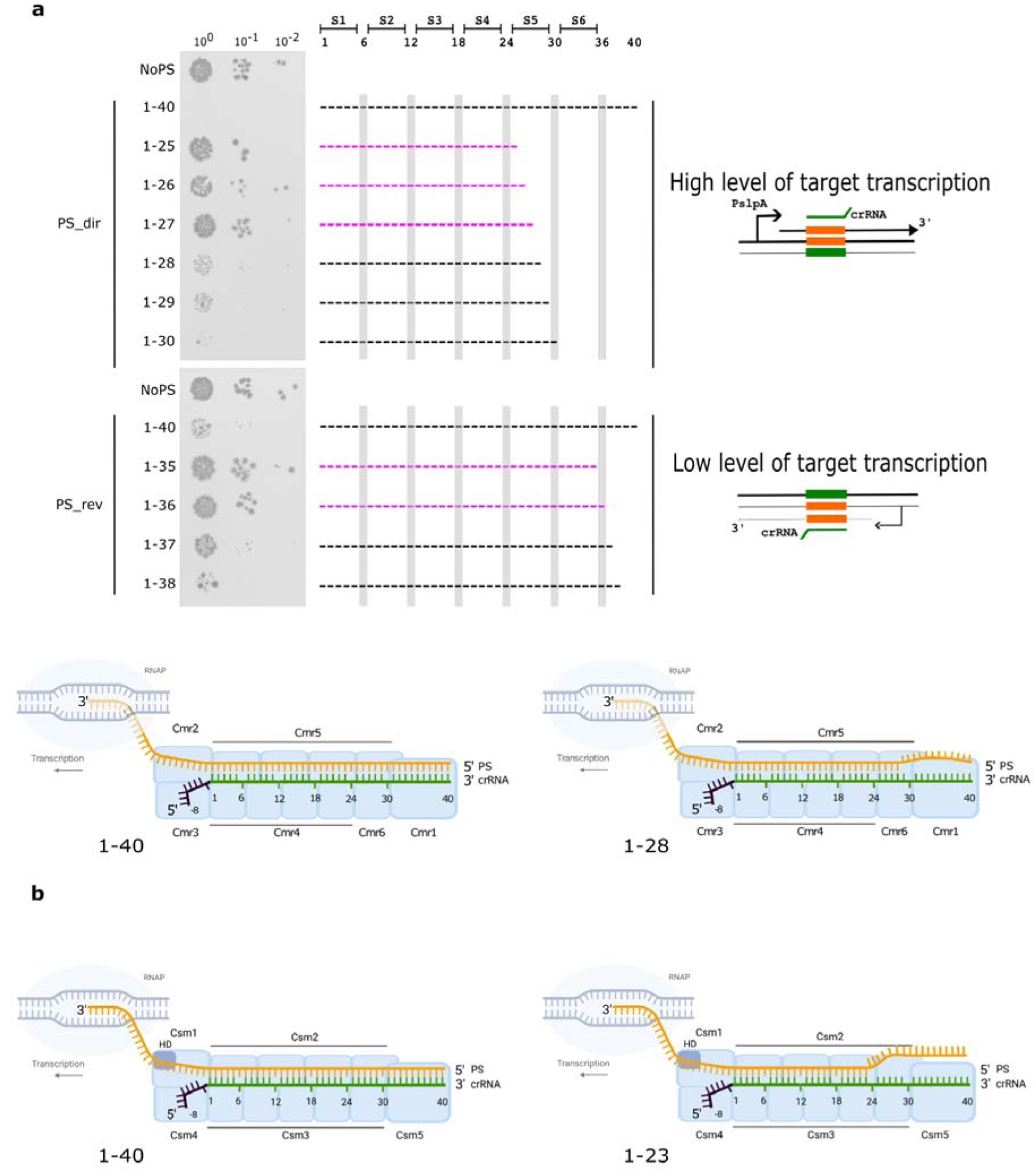
Defining spacer-protospacer complementarity requirements at the 3’ end-proximal part of the crRNA spacer for *T. thermophilus* Type III-B interference with plasmid transformation. **a.** Results of transformation of PS_dir and PS_rev derivatives with nested multiple mismatches with the 3’ end part of the crRNA spacer into the III-B^+^ strain are shown. See Fig. 1 legend for details. Results of a similar experiment with the III-A^+^ strain can be found in You et al. [54]. **b.** Schematic structures of effector complexes with a fully matching target (left) and with maximally mismatched highly transcribed targets that support interference (right) are shown. For the III-A effector, the data are taken from You et al. [54].

The differences in target recognition requirements by the III-A and III-B systems could be caused by different amounts of effectors present in the cells. To determine abundance of III-A and III-B effector subunits, we analyzed the *T. thermophilus* HB27 proteome using liquid chromatography coupled to tandem mass spectrometry (LC-MS/MS). To compare the relative amounts of effector subunits, their intensity based absolute quantitation (iBAQ) signals were normalized on the total protein iBAQ signal. The iBAQ signal of the RNA polymerase alpha subunit was used as a reference. In fast growing bacteria there are several thousand RNA polymerase molecules per cell [57]. In Class I effectors, the stoichiometry of Cas7-like subunits (Csm3 for III-A and Cmr4 for III-B) is the highest [25]. Consistently, the iBAQ signals from these subunits were the strongest. While Csm3 was detected in each of the nine replicates, the Cmr4 signal was detected only in three. The mean signal from Csm3 was 10 times higher than that for Cmr4 (Supplementary Figure S4a). In turn, the signal from Csm3 was ∼10 less than from the RNA polymerase subunit. Our analysis also revealed iBAQ signals for Cas7 subunits of the I-B and I-C effectors, which were comparable to the Csm3 signal (Supplementary Figure S4a). Overall, we conclude that the III-A effector subunits are presented in much larger quantities than their III-B homologs, which may be the reason for observed weaker III-B interference against less transcribed targets and more stringent target complementarity requirements for this system.

### Determining the minimal region of crRNA-target complementarity sufficient for III-A interference

We next combined protospacer mutations introducing mismatches with the 3’ and 5’-end proximal parts of the crRNA spacer and determined the transformation efficiency of plasmids harboring mutated protospacers. Only the III-A^+^ strain and PS_dir variants were tested, as this combination gave the strongest interference, was least sensitive to mismatches between the target and crRNA, and no small transformant colonies that complicate data interpretation were observed. We began with the PS_dir variant mismatched with crRNA after position +26 (subject to interference, see [54] and introduced additional mismatches in positions +1 and +1/+2. The single mismatch had no effect on interference, while the double one abolished it (Fig. 3), indicating that with this target the minimal crRNA-target duplex sufficient for interference spans positions +2/+26. To determine whether the position of this minimal duplex within the effector is important for interference, we combined mutations introducing 5’ end proximal mismatches with mutations that introduced mismatches after position +27. While a mismatch extending to +2 abolished interference on the background of mismatches past position +28, it had no effect when combined with a mismatch past position +29 (a +3/+28 duplex). A mismatch extending to +3 abolished interference with a mismatch past position +29 but supported interference on the background of a target mismatched after position +30 (a +4/+29 duplex). A mismatch extending to position +4 abolished interference on the background of a target mismatched after position +30 but was interference-proficient when complementarity with the target was extended to position +31 (a +5/+30 duplex). The pattern did not repeat itself with a mismatch extending to position +5, which was not subject to interference on the background of duplexes ending either at +31 or +32. However, for the mismatch extending to +6, interference was observed in combination with a mismatch after position +33 (a duplex of +7/+32) but not after position +32 (a duplex of +7/+31). Neither the +8/+32 nor +8/+33 duplex was functional, however. Extension of the duplex boundary beyond the position +9 was not attempted, as it did not support interference even in the absence of mismatches with the 3’ end proximal part of crRNA (Fig. 1a). Overall, we conclude that for III-A interference against a highly transcribed target, there has to be a region of complementarity between crRNA and the target that extends at least over 24/25 base pairs. This region can be shifted downstream by 6 base pairs from position +1. While this has not been tested, we assume the same would hold true for III-A interference against PS_rev, but the area of complementarity sufficient for interference would be more extended. We also assume that similar trends will hold true for the III-B interference.

**Figure 3.**
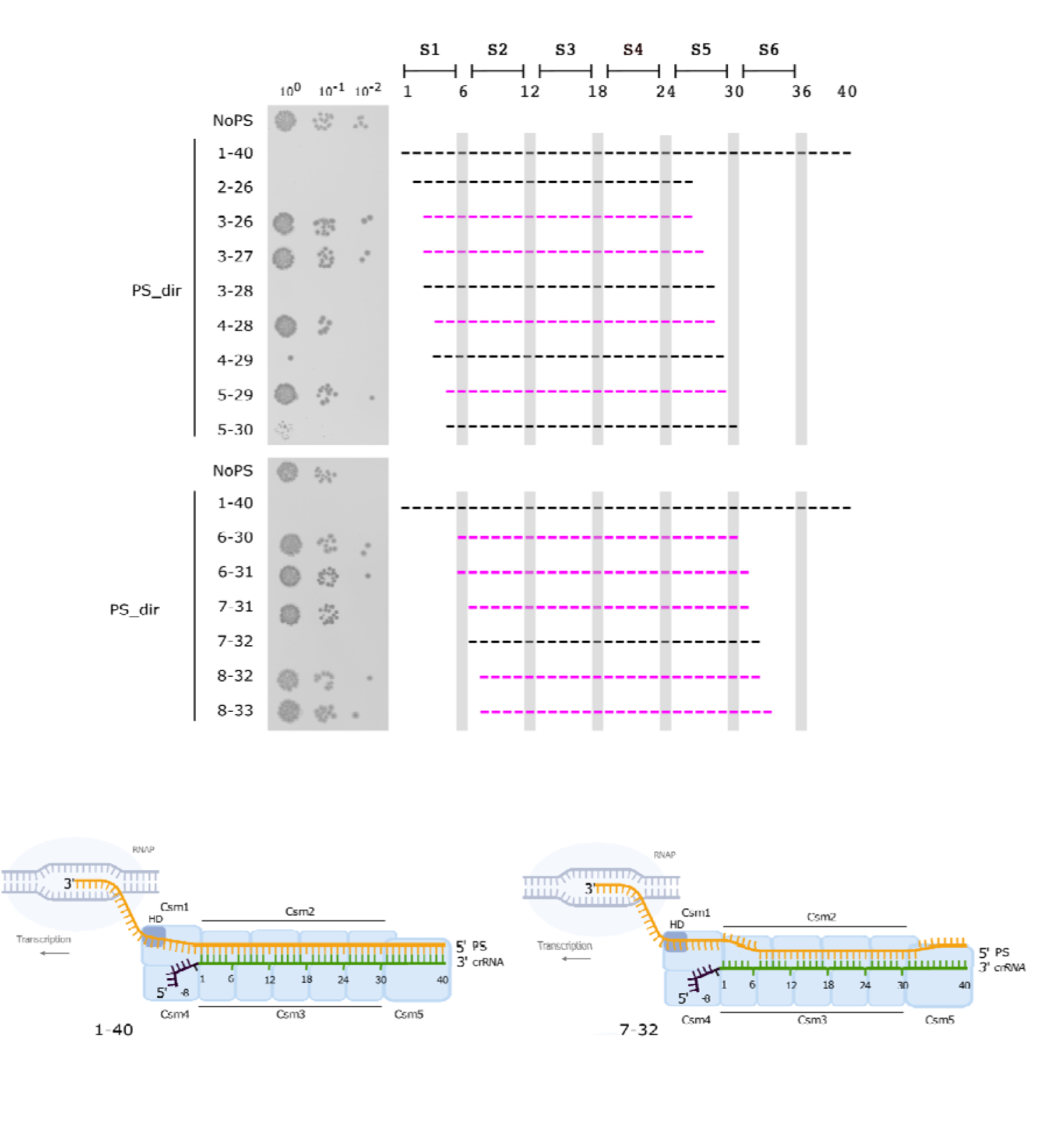
Defining the minimal length of continuous RNA duplex sufficient for *T. thermophilus* Type III-A interference with plasmid transformation. Results for PS_dir plasmid and its derivatives are presented. See Fig. 1 legend for details.

### Interrupted crRNA-target duplexes allow interference

We next introduced internal mismatched segments between crRNA and the PS_dir target and determined their effects on III-A and III-B interference. Mismatched segments were 5 nucleotides long and each corresponded to a segment of crRNA that is bound by one Csm3 (III-A) or Cmr4 (III-B) subunit. Data presented above already show that prevention of duplex formation by the part of crRNA that interacts with the first Csm3 subunit (S1), does not affect III-A interference. Mismatches in segments bound by the second, third, fourth, and fifth Csm3 subunit similarly had no or partial (most notably, for the S2 segment) effect on interference (Fig. 4a). For the III-B^+^ strain, only disruption of base pairing in the S4 segment was tolerated (Fig. 4b). Thus, continuous duplex is not required for Type III interference in *T. thermophilus*. We note that our results for III-B immunity are in contrast with earlier *in vitro* findings of [37] who reported that the +19/+23 (S4) mismatch abolished cleavage of target RNA.

**Figure 4.**
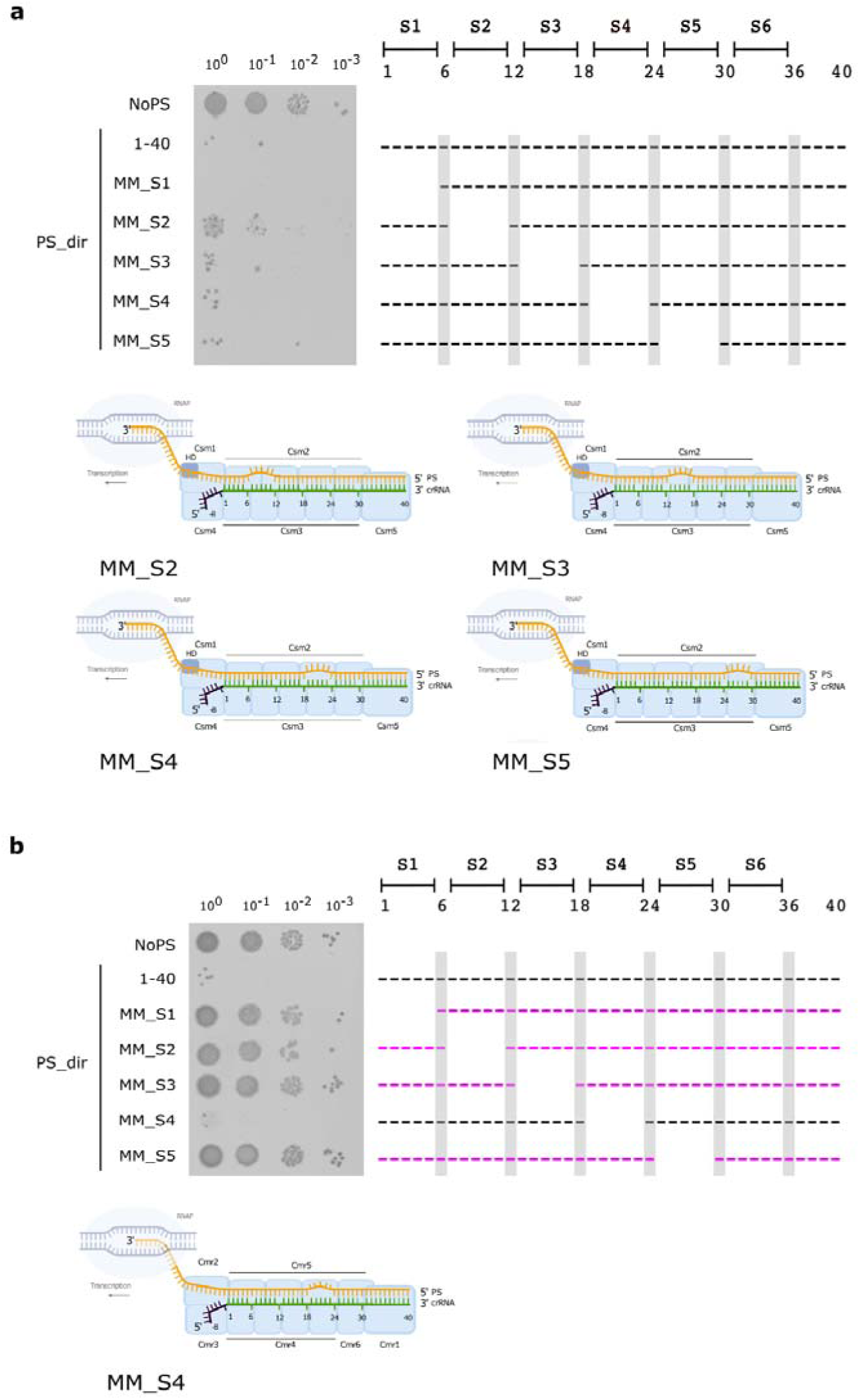
Defining the impact of internal duplex segments mismatches on *T. thermophilus* Type III-A (a) and III-B (b) interference. Results for the PS_dir plasmid and its derivatives are presented. See Fig. 1 legend for details. The schemes at the bottom of each panel show interference-proficient effector-target complexes.

### Impact of spacer-protospacer mismatches by the crRNA 5’ handle

An interaction of the III-A effector with a target mismatched at positions +1/+7 must result in a complex that contains a single-stranded piece of crRNA between the duplex and the 5’ crRNA handle, which is bound to the Csm4 subunit. Base pairing at the handle region with the target prevents interference [22]. To test whether control by handle base pairing is still operational with a shortened duplex, we tested the effect of mutations in PS_dir that made the handle complementary to RNA upstream of the protospacer. In the case of fully matching spacer-protospacer pair, extension of base pairing to the handle region abolished interference, as expected (Fig. 5). The same effect was observed on the background of the +1/+7 mismatch, indicating that the handle can base pair with the target and switches off interference even when the first Csm3 subunit is not interacting with the target.

**Figure 5.**
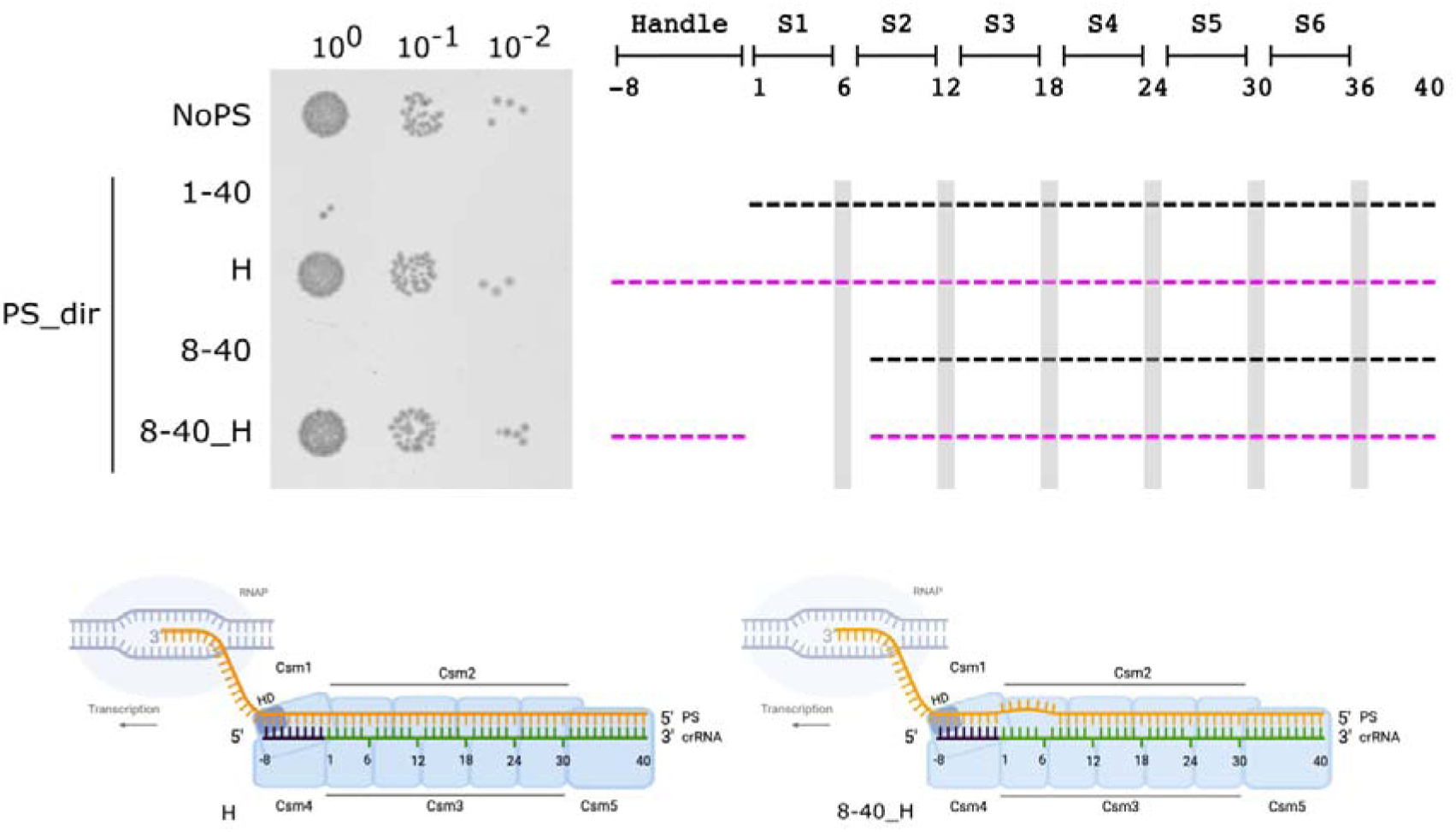
Complementarity of protospacer transcript with crRNA handle abolishes Type III-A interference when the first 7 positions of the spacer are mismatched. Testing the III-A^+^ strain transformation efficiency by indicated PS_dir derivatives. In derivatives carrying the “H” label, the 5’-handle of crRNA is complementary to the target. See Fig. 1 legend for details. Schematic representation of non-active Type III effector complex with the crRNA 5’ handle complementary to the target is shown at the bottom (“H”, left). The likely structure of a similar complex with mismatched crRNA-target positions +1/+7 is shown on the right.

### The role of the Cas10 HD domain and cOA-sensing CARF domain containing nucleases in Type III-A immunity

The HD nuclease domain of Type III effectors’ Cas10 subunit is responsible for target-activated DNA cleavage that may contribute to interference with plasmid transformation [34]. We created a III-A^+^ strain where the catalytic histidine (H) and aspartate (D) residues of the HD domain were replaced for alanines. As judged by plasmid transformation efficiency, the HD mutant no longer interfered with either PS_dir or PS_rev, indicating that the HD nuclease is necessary for Type III-A immunity. However, compared to the pMK18 vector control, PS_dir (but not PS_rev) colonies formed by transformed HD mutant cells were markedly smaller (Fig. 6a), suggesting slower growth. Presumably, this growth inhibition is due to cOAs synthesized by the target bound III-A effector. The growth inhibition effect was also observed in HD mutant colonies transformed with a PS_dir derivative mismatched up to position +7, but not with a derivative mismatched up to position +8 that is not subject to III-A interference (see above).

**Figure 6.**
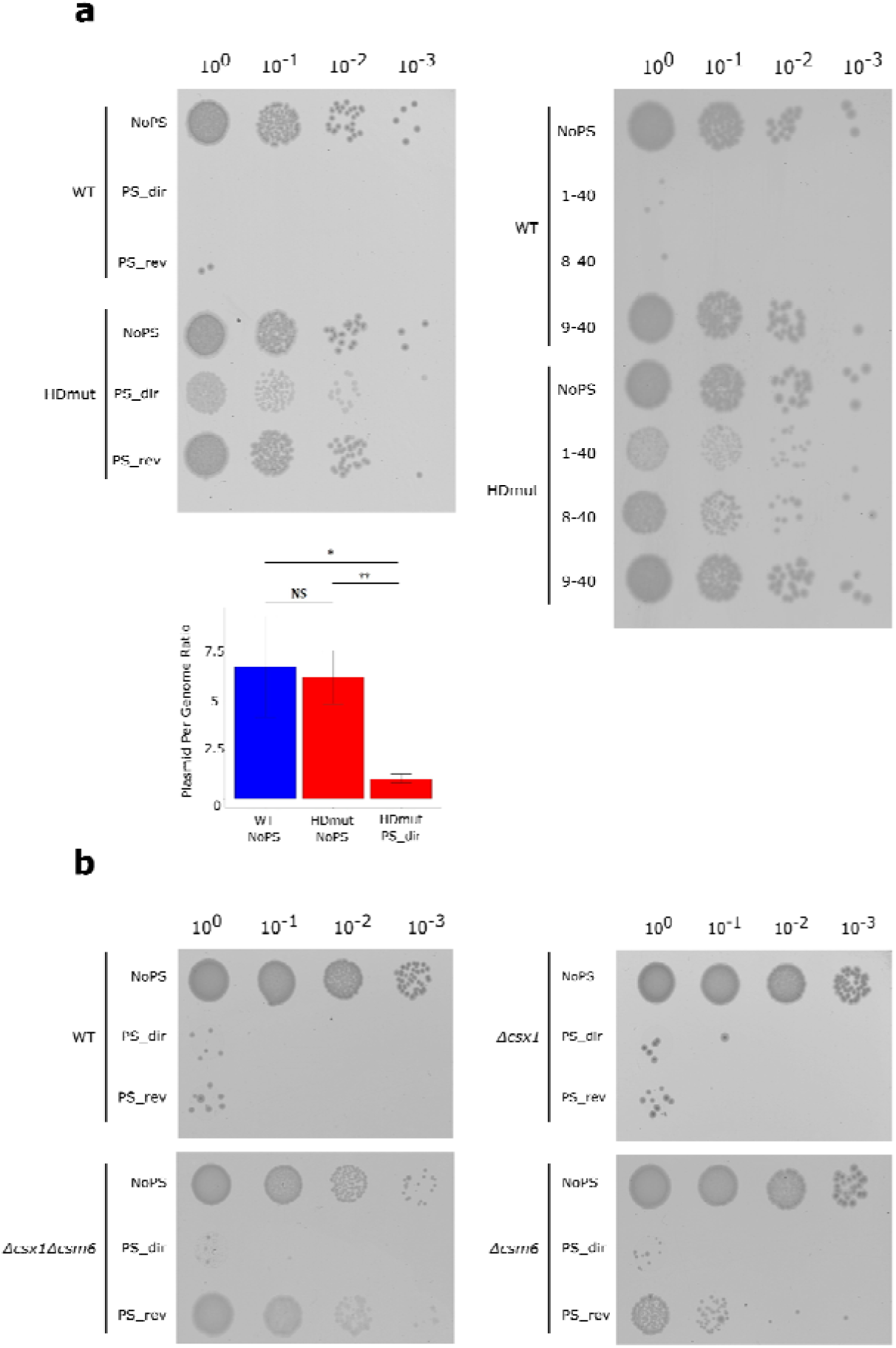
Effects of Cas10 HD domain (a) and cOA-sensing CARF domain containing nucleases (b) mutations on Type III interference. **a.** Testing the transformation efficiency by indicated PS_dir derivatives into the III-A^+^ strain (wt) or III-A^+^ mutant with disrupted Cas10 subunit HD domain “HDmut”. See Fig. 1 legend for details. At the bottom, plasmid per genome ratio in indicated transformed colonies is shown. **b.** Testing the transformation efficiency of PS_dir, PS_rev, and the pMK18 control (“NoPS”) into wild-type *T. thermophilus* HB27 or mutants lacking the indicated genes coding for CARF domain containing nuclease.

We hypothesized that the slower growth of HD mutant cells carrying the target plasmid on selective medium is due to decreased plasmid copy number. To test this idea, we performed the qPCR assay with a pair of primers specific for pMK18 plasmids and another one, specific for the chromosomal *rho* gene. The results indicated that compared to wild-type cells, plasmid copy number was decreased 3-4 fold in cells with the inactive HD domain (Fig. 6a) [56]. We conclude that the Cas10 HD domain plays an important function in *T. thermophilus* III-A immunity. Without this domain activity, the III-A system is not able to mount efficient interference: though target recognition still occurs, plasmids carrying the target cannot be completely cleared from the cell.

Since the data obtained with the HD mutant indicate that partial Type III interference can be observed when some arms of the defense mechanism are switched off, we investigated the role of *T. thermophilus* cOA-sensing CARF domain containing nucleases Csx1, Csm6, and Can1. To this end, we attempted to knock out each of the three genes or their combinations. While we encountered technical difficulties during the construction of the Δ*can1* mutant, single Δ*csx1* and Δ*csm6* mutants, as well as the corresponding double mutant were prepared. Because of the limited number of selective markers available for genomic engineering in *T. thermophilus*, the mutants were constructed on the wild-type background containing functional III-A and III-B systems. Transformation with pMK18 control and PS_dir and PS_rev plasmids showed that the Δ*csx1* mutant behaved as the wild-type and was poorly transformed with either PS_dir or PS_rev (Fig. 6b). The Δ*csm6* mutant efficiently interfered with PS_dir but produced ∼10 times more transformants with PS_rev, indicating that Csm6 activity makes interference with targets of low abundance more efficient. The transformant colonies were smaller than colonies transformed with pMK18, possibly suggesting decreased plasmid copy number. The double mutant interfered with PS_dir as efficiently as wild-type, but interference against PS_rev was either absent or very low.

We employed LC-MS/MS to determine the abundance of CARF domain containing nucleases Csx1, Csm6, and Can1. The Csm6 signal was consistently detected in all replicates and was comparable to the Csm3 signal (Supplementary Fig. S4b). Signals from Csx1 and Can1 were not detected. Thus, while Csm6 is clearly involved in Type III immunity, Csx1 and Can1 may contribute when their intracellular amounts are increased due to specific environmental/physiological cues.

## Discussion

Effects of mismatches between crRNAs and their target for Type III CRISPR-Cas interference have been studied both *in vitro* and *in vivo*. However, interpretation of the data has been complicated. *In vivo*, conflicting results were obtained with different experimental models used to monitor interference (i.e., plasmid transformation or conjugation versus phage infection). Since Type III interference efficiency depends on the abundance of target RNA, the results may depend on both the expression level and the time when the targeted transcript is expressed, further confounding comparison of different sets of data [58]. Given that Type III systems interfere with phage infection and plasmid transformation even in cases when the target transcripts are not required for the infection/plasmid maintenance, the Type III immune response must be mediated by the activities of target-bound effector complexes (presumably, by the cyclase activity of the Cas10 subunit and, when present, by the nuclease activity of its HD domain). The dependence on target concentration implies, in turn, that the strength of various interference-related responses should depend on the concentration of effector species loaded with a particular crRNA, which may explain apparent differences between spacer-protospacer pairs reported in *in vitro* studies with effectors loaded with multiple crRNAs purified from cells [37].

In this work, we studied target complementarity requirements of two *T. thermophilus* Type III subtypes by monitoring plasmid transformation efficiency. Our experimental system allowed us to compare interference requirements for identical targets of different abundance by each Type III effector subtype. We found that both III-A and III-B systems, alone, decreased formation of colonies by cells transformed with a plasmid producing highly abundant transcript by at least 100-fold. Colonies that did appear were escapers. Approximately half of escaper colonies were formed by cells carrying point mutations in Type III *cas* genes essential for interference. Another half belonged to an interesting and, to our knowledge, novel class. They resulted from recombination between identically oriented Type III arrays located at both sides of deleted *cas* operons (Supplementary Fig. S2). Escaper-generating recombination events produced new hybrid arrays at the sites of deletion ends junctions. Closer inspection revealed that recombination involved repeats located throughout the length of both arrays (Supplementary Fig. S3). Thus, CRISPR repeats are recombinogenic, which may provide a mechanism for mobility of CRISPR-Cas systems when *cas* operons are flanked by arrays in the same orientation. Our data suggest that recombining arrays can be quite far away from each other. For example, the largest deletions observed in escapers removed ∼90 kbp from the 256 kbp megaplasmid pTT27 that encodes both Type III *cas* operons.

In agreement with earlier data [54], we found that crRNA-target complementarity requirements for III-A and III-B interference depend on the level of target abundance. A ca. two orders of magnitude decrease in target abundance increases the minimal length of crRNA-target duplex required for interference at the 3’ end of the duplex by 3 base pairs for the Type III-A effector (from 1-23 to 1-26) and by 8 base pairs (from 1-30 to 1-38) for the Type III-B effector. The 5’ end of the duplex that supports interference starts at positions +7 or +4 for, correspondingly, high and low abundance targets for the Type III-A effector and positions +4 and +1 for the Type III-B one. The observed dependence of complementarity requirements on target abundance suggests that for Type III effectors target recognition is driven by the strength of crRNA-target interactions, and that the onset of interference occurs when the concentration of target-bound effector complexes exceeds a certain threshold. We used a simple thermodynamic argument to understand the dependence of Type III interference on the abundance of target RNA and the minimal duplex length. We consider reversible binding of *effector* to target transcript (*TT*): (1)

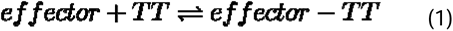

and assume that the stationary concentration of the effector-target complex [*effector – TT*] is determined by the mass action equilibrium:

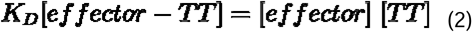

The dissociation constant *K_D_* is expressed as the exponent of the Gibbs free energy difference of the reactants in (1):

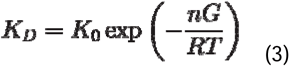

Here, *K_0_* comprises a dimensional prefactor and the exponent of the Gibbs free energy of “residual” binding of the CRISPR machinery to target RNA that does not depend on nucleotide complementarity. We assume that the free energy of binding of a pair of complementary nucleotides Δ*G* is additive and identical for all complementary base pairs. The number of complementary nucleotides is denoted as *n* and non-complementary bases are considered as non-interacting. We also assume that the concentration of effector-target RNA complex is much smaller than the free concentrations of effector bound to a particular crRNA and the target RNA concentration. In other words, [*effector*] and [*TT*] in the right hand side of the equilibrium are understood as total concentrations.

Consider two cases that differ by the number of complementary nucleotides and concentrations of target RNA [*TT*]:

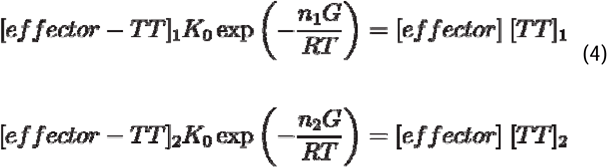

We assume that the concentration of effector-target RNA complex that is necessary and sufficient to induce the interference response (through activation of cOAs synthesis and/or HD domain nuclease) is the same in both cases, i.e., [*effector – TT*]_1_ = [*effector – TT*]_2_. To determine how the difference in concentrations of target, (parametrized here as their ratio [*TT*_1_]/[*TT*_2_]), translates into a difference in the number of complementary base pairs between crRNA and the target *n*_2_ – *n*_1_ necessary for interference, we divide the first line of (4) by the second one, taking into account that [*effector – TT*]_1_ = [*effector – TT*]_2_

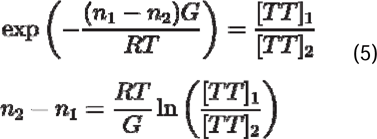

Numerical evaluation of (5) requires the knowledge of Δ*G* = Δ*H* – *T*Δ*S* (enthalpy and entropy of binding), averaged over all complementary base pairs. Based on the data of Ghosh et al. [59], these averages are Δ*H ≈* –11.7*kcal/mol* and Δ*S* ≈ –31.6 cal/mol°*K*.

Evaluating the equations for a particular example of target requirements for *T. thermophilus* Type III-A interference with *T* = 338°*K*, and [*TT*]_1_/[*TT*]_2_ = 80 yields *n*_2_ – *n*_1_ ≈ 2.9, which is an almost perfect match with the experimental value of (*n*_2_ – *n*_1_)*_exp_* = 3. The fact that the difference in complementarity requirements with the 3’-end proximal part of crRNA in the case of III-B effector is increased to 8 base pairs suggests that the contribution of this part to target binding is less than that of the 5’ end proximal part, which agrees with our experimental observations that disruption of complementarity between longer segments of 3’-end proximal part of crRNA and the target is tolerated.

Shortening of the crRNA-target duplex at both sides revealed that Type III-A interference against a highly abundant target is equally effective when effector forms heteroduplexes corresponding to positions 2-26, 3-28, 4-29, 5-30, and 7-32. Further shortening of the duplex at either end abolished interference. Thus, it appears that the Type III-A effector can bind the target forming a minimal ∼24-25 bp duplex and the position of the duplex within the effector is flexible. While this has not been tested, we assume the same would hold true for Type III-A interference against PS_rev, but the area of complementarity sufficient for interference would become extended. We also assume that similar trends will hold true for the Type III-B interference.

In our assays, Type III interference reveals itself by the absence of colonies of cells carrying protospacer-containing plasmids. Since protospacer transcripts are not essential for plasmid maintenance, it follows that the lack of transformants must be due to *i*) destruction of plasmids from which nascent target transcripts are transcribed, *ii*) suicide or significant growth rate reduction of cells in which a Type III target is recognized. Co-transcriptional destruction of DNA by a Type III effector has been demonstrated *in vitro* in a pure system [60] (though later disputed [61]) and would require a nuclease activity provided by the HD domain of the Cas10 subunit in Type III-A or by the cOA-activated protein Can1 in both Type III-A and III-B systems. Indeed, a III-A^+^ strain carrying inactivating substitutions in the HD domain catalytic residues is transformed efficiently, but the colonies formed are small, indicating growth inhibition caused by the recognition of target RNA by the effector, since cells harboring a plasmid with a mutated target that is not subject to interference are forming normal, healthy colonies. The plasmid copy number in cells from small colonies is decreased, consistent with a view of an ongoing interference that is too weak to completely purge the plasmid. The lower copy number may be sufficient to decrease growth on antibiotic-supplemented medium that selects for plasmid presence. The slow growth may also be caused by poisoning of cells by secondary effects induced by target recognition [34].

The fact that *T. thermophilus* encodes two functional Type III CRISPR-Cas systems employing shared CRISPR arrays raises a question regarding the potential adaptive value of this apparent redundancy. Although in our experiments the III-A^+^ and III-B^+^ strains interfere with the same efficiency against highly transcribed targets, one can imagine that in the case of some invaders activities of both systems will be necessary for optimal defense. This could be due to non-trivial mechanistic differences between the Type III-A and III-B immunity. The current view of the Type III defense mechanism implies three arms of the immune response (target transcript degradation, followed by target-dependent DNA cleavage, and cOA-activated nonspecific RNA cleavage). Recent systematic analysis shows that more than half of Type III-B Cas10 proteins lack the HD domain while many others have it inactivated [62], which implies the lack of transcription-dependent DNase activity. Whatever the target of this activity, our data as well as the data by others show that it is necessary for optimal Type III-A immunity. Interestingly, in the absence of active HD domain, the level of protection provided by *T. thermophilus* Type III-A system is less than that provided by intact Type III-B system, despite the fact that the III-B effector is much less abundant. One can therefore speculate that the Type III-B defense relies on additional pathways that are not involved in the Type III-A defense. At first glance, this hypothetical pathway(s) should not be cOA-dependent, as both the Type III-A and III-B effectors produce cOAs. However, it is possible that the largest subunits of Type III-A and III-B effectors produce not fully identical sets of oligonucleotides, which results in activation of distinct CARF-domain nucleases. Consistently, a great variety of cOA-activated effectors specifically associated with Type III-B loci was recently discovered [31,40,63,64]. Another plausible explanation for the observed discrepancies between the immunity responses maintained by Type III-A and III-B systems relies on the different kinetics of the reactions catalyzed by the effectors. Differences in the compositions of the effector complexes and potentially varying rates of target RNA cleavage and dissociation of the cleavage products could lead to distinct durations of the activated states of the effectors. This is significant because the cleavage of target RNA with the subsequent dissociation of the products switches off the cyclase activity of Cas10 subunits, consequently switching off downstream cOA-activated mechanisms [65,66]. In the scenario where the III-B effector-target complex is more stable it could potentially result in a more efficient cOA-dependent response. In agreement with the suggestion, recent experiments on the quantification of the cOAs produced by Type III-A and III-B effectors of *T. thermophilus in vitro* shows that, in the same experimental conditions, Type III-B effectors produce ca. 3 times more cOAs than Type III-A effectors [67]. One can further speculate that by keeping two subtypes of Type III systems of which one mostly relies on cOA-activated effectors and another - on HD domain nucleases, cells are able to better cope with mobile genetic elements that employ anti-CRISPR proteins targeting just one arm of the defense.

Since we did not detect a region that could be considered as a *sensu stricto* “seed” required for target transcript recognition, a question about the mechanism of efficient target transcript search by the Type III effectors arises. Taylor et al. noticed that crRNA base flipping mediated by Cmr4 (and, therefore, also by Csm3) subunits in the assembled effector complex filament introduce periodical disruptions in the duplex between the target and spacer part of crRNA that resemble DNA-DNA interactions mediated by RecA [52,68]. During the homology search, segments of RecA filaments may probe the target DNA independently, which should accelerate the search process [69]. A common mechanism may be employed during target search by Type III effectors, where at least some targets can be highly structured because of secondary and tertiary interactions in RNA. Our data show that disruptions of base pairing by crRNA segments bound by individual Csm3, including those located inside the full-length RNA duplex, have no or partial effect on the more efficient *T. thermophilus* III-A interference. This is consistent with independent recognition of 5 nucleotides long target segments by the effector. Nucleation of duplex formation by multiple segments of crRNA thus allows target recognition without a seed.

## Materials and methods

### Strains and cultivation conditions

*Escherichia coli* DH5α (F− Φ80lacZΔM15 Δ(lacZYAargF) U169 recA1 endA1 hsdR17(r_k_^−^, m_k_^+^) phoA supE44 thi-1 gyrA96 relA1 λ−) was used for molecular cloning and was cultivated in LB medium at 37°C. Agar was added up to 1.5% (w/v) for growth on plates. *T. thermophilus HB27* cells were cultivated in TBM medium (0.8% w/v tryptone, 0.4% w/v NaCl, 0.2% w/v yeast extract in Vittel mineral water) supplemented with 0.5 mM MgSO_4_ and 0.5 mM CaCl_2_ at 70°C and 180 rpm in an Environmental Shaker-Incubator ES-20/60 (Biosan). Agar was added up to 2% (w/v) for growth on plates. Plates were incubated at 65°C in a RedLine RI 53 (Binder) incubator.

*T. thermophilus* strains with deactivated III-B (Δ*cmr4*) or III-A (Δ*csm3*) interference modules were previously constructed in our laboratory [54]. *T. thermophilus* Δ*csx1*, Δ*csm6* and Δ*csx1*Δ*csm6* mutants were constructed by replacing corresponding genes of wild-type *T*. *thermophilus* HB27 with thermostable antibiotic resistance markers through natural homologous recombination following the procedure described in [53].

The III-A^+^ strain derivative encoding the *csm1* gene with catalytically inactive HD domain (“HDmut”) was prepared by gene editing through homologous recombination using the pT7_bleoHDmut recombination plasmid. To create pT7_bleoHDmut, the bleomycin resistance cassette was amplified from plasmid pWUR112 [70] and the recombining flanking regions were amplified from *T. thermophilus* HB27 genomic DNA. These fragments were assembled into the pT7blue plasmid using Gibson Assembly Mix and transformed into *E. coli* DH5α cells, and plated on LB agar plates supplemented with 100 μg/ml ampicillin. Clones of interest were selected by PCR and plasmids from selected clones were purified and verified by Sanger sequencing. Next, point mutations were introduced using appropriate primers. The mutated product was cloned back into the pT7blue plasmid by restriction/ligation cloning, transformed into *E. coli* DH5α cells, and plated on LB-agar plates supplemented with 100 μg/ml ampicillin. Clones of interest were selected by PCR and plasmids from selected clones were purified and verified by Sanger sequencing.

III-A^+^ cells were transformed with the pT7Blue-derived plasmid encoding bleomycin resistance cassette [70] flanked with nucleotide sequences homologous to the 5’-proximal part of the *csm1* gene and the genomic sequence upstream the 5’ end of the *csm1* gene. The flanking site homologous to the 5’-proximal part of the *csm1* gene was constructed in such a way that, in case of homologous recombination, nucleotides encoding the catalytic histidine and aspartate residues of the Csm1 HD domain will be replaced with nucleotides encoding two alanine residues.

*T. thermophilus* cells were transformed with the pT7_bleoHDmut and plated on TBM-agar plates supplemented with 30 μg/ml bleomycin. After 3 passages on selective media, the identity of selected clones was confirmed by PCR and whole-genome sequencing. To account for potential effects of the bleomycin resistance cassette integrated into the type III-A locus, a strain containing the same resistance cassette upstream of intact *csm1* gene was prepared in a similar way and used as control in interference assays.

### CRISPR Interference Assay

pMK18-based plasmids carrying protospacers with mismatched spacer segments were constructed as described previously [54] using appropriate double-stranded oligonucleotides (synthesized by Evrogen, Russia, listed in Supplementary Table S1). To obtain mismatched positions, protospacer nucleotides were changed to their complementary base pairs. Kanamycin (50 μg/ml for *E. coli* and 30 μg/ml for *T. thermophilus*) was used to select pMK18-containing bacteria. *T. thermophilus* cells were transformed with pMK18 or protospacer-bearing derivatives according to the protocol described previously [56]. Serial 10-fold dilutions of the transformation mixtures were prepared, and 10 μL drops of the diluted transformation mixtures were deposited on the surface of TBM agar plates containing kanamycin, and results were recorded after growth for 18-42 hours.

### Extraction of Genomic DNA

1-2 ml of overnight *T. thermophilus* HB27 cultures was centrifuged at 3000 g for 2 min. Cell pellet was resuspended in 250 μl of TES buffer (10 mM Tris-HCl (pH 8.0), 1 mM EDTA, 0.5% SDS, 20 μg/ml of Proteinase K) and incubated at 56°C for 30 min. The sample was extracted with 0.5 ml of phenol (pH 8.0) at 60°C for 20 min. The solution was then chilled and centrifuged at 12°C for 15 minutes. The aqueous phase was extracted twice with 200 μl of chloroform. Nucleic acids from the aqueous phase were ethanol precipitated at -70°C in the presence of 1/10 volume of 3 M sodium acetate (pH 5.2). After centrifugation, the resulting pellet was washed twice with 70% cold ethanol, once with 96% cold ethanol, dried at room temperature, and resuspended in either TE buffer (10 mM Tris–HCl (pH 8.0), 1 mM EDTA) or Milli-Q water.

### Extraction of total RNA and high-throughput RNA sequencing

The procedure of total RNA extraction, library preparation and sequencing is described in previous work [54]. Reads were processed and mapped onto reference sequences (*T. thermophilus* HB27 genome assembly GCF_000008125.1 and pMK18 plasmid) using the READemption tool with default parameters [71]. Alignment files were used to estimate the abundances of transcripts mapped onto the either strand of the pMK18 polylinker site. Due to the short length of the polylinker site, a larger region downstream *kanR* gene was selected for the analysis to decrease effects related to unevenness of the coverage. The number of alignments intersecting with each strand of the selected region were calculated using the Rsubread package [72]. The transcript per million (TPM) values were calculated in R, and the plots were prepared using custom scripts.

### Whole-genome sequencing

Individual III-A^+^ and III-B^+^ colonies (10 each) formed after PS_dir transformation were picked at random, transferred into 5 ml of TBM supplemented with kanamycin and grown overnight. DNA was extracted and subjected to high-throughput sequencing on the Illumina NextSeq platform. DNA libraries were prepared with NEBNext Ultra II kit (New England Biolabs). For clones bearing extensive deletions, sequencing was additionally performed using Oxford Nanopore Technologies (ONT). DNA libraries were prepared from the non-sheared DNA using the Native Barcoding kit (SQK-NBD114-24, ONT, Oxford, UK) with enrichment of long fragments using the Long Fragment Buffer (LFB) according to the manufacturer’s instructions (ONT, Oxford, UK). Sequencing was performed on MinION using the R10.4.1 flow cell (FLO-MIN114, ONT, Oxford, UK) with a translocation rate of 400 bps. Basecalling was carried out using Guppy 6.0.1 in the High accuracy model.

The adapter removing, quality trimming and filtering procedures were performed using Trimmomatic v0.39 [73]. The procedures of reads mapping, variant calling and functional annotation of detected variants were performed by snippy pipeline [74]. The obtained alignments and variants were visualized using IGV browser [75]. The hybrid genome assemblies from both short paired Illumina reads and long Oxford Nanopore reads were prepared using Unicycler pipeline [76].

### Analysis of the recombination events in the escapers

To analyze recombination events affecting CRISPR arrays in III-A^+^ and III-B^+^ strains, colonies obtained on kanamycin-containing TBM agar after transformation with PS_dir pooled. Each plate contained ∼500 colonies and three transformations were used for each strain. DNA was purified and DNA fragments corresponding to expected products of recombination events were amplified using Phusion High-Fidelity Polymerase (Thermo Fisher Scientific, USA). Standard PCR running conditions were used with three primer pairs flanking the CRISPR-5 and CRISPR-2 (Th_RecCRISPR-F2 and Th_RecCRISPR-R2 primers), CRISPR-5 and CRISPR-11 (Th_RecCRISPR-F1 and Th_RecCRISPR-R2 primers), and CRISPR-2 and CRISPR-11 (Th_RecCRISPR-F1 and Th_RecCRISPR-R1 primers, all primers are listed in Supplementary Table S1). Amplification products were visualized by agarose gel electrophoresis and purified from the gel. Amplicons were then subjected to the sequencing on the Oxford Nanopore MinION platform using the R10.4.1 flow cell (FLO-MIN114; ONT, Oxford, UK) with a translocation rate of 400 bps. Sequencing libraries were prepared with Native Barcoding kit (SQK-NBD114-24, ONT, Oxford, UK) with enrichment of short fragments using the Short Fragment Buffer (SFB) according to the manufacturer’s instructions (ONT, Oxford, UK). Basecalling was carried out using Guppy 6.0.1 in the High accuracy model. Oxford Nanopore reads containing nucleotide sequences of primers used for amplification were selected using custom scripts. Search of CRISPR arrays in the selected reads was performed by CRISPRCasFinder tool [77]. To determine the origin of spacers in detected arrays, sequences of the spacers were extracted from reads and compared to the set of spacers in the *T. thermophilus* HB27 genome using the blastn tool [78]. The CRISPR array was considered as a product of the recombination event between two CRISPR arrays if the analyzed array 1) contained the spacers derived from both parental arrays, 2) the order of the spacers matched to the parental array, coincided with the order of spacers in the parental array. For final processing of the data and plot construction, custom R and Python were used.

### Determination of plasmid and chromosomal DNA concentration

Cells were grown in TB media until OD_600_ = 0.6, harvested by centrifugation and total DNA was extracted as described above. qPCR analysis of *rho* (genome DNA) and *kanR* (plasmid DNA) genes was carried out using Maxima SYBR Green/ROX qPCR Master Mix (Thermo Fisher Scientific) and Rho_F/Rho_R and Kan_F/Kan_R primer pairs, correspondingly. Primer efficiencies for both primer pairs were obtained using assays with serial dilutions of a mixture of genomic DNA and pMK18 with known concentrations. Plasmid per genome ratio was calculated based on normalized (with primer efficiency) ratio of *kanR* and *rho* qPCR signals.

### Proteomic analysis

To prepare trypsin cell hydrolysates, cell pellets were resuspended in 10 mM Tris-HCl (pH 7.0), 1 mM phenylmethylsulfonyl fluoride (PMSF) and disrupted by sonication. After centrifugation (10 min, 15,000 g, 4°C), an equal volume of methanol-chloroform (1:0.75) was added to cleared lysates. After phase separation, proteins from the interphase were collected, dried at 50°C, resuspended in a denaturation buffer (10 mM Tris-HCl (pH 8.0), 6 M urea, 2 M thiourea), heated at 95°C for 5 min. For cysteine reduction, 1 mM DTT was added. The reactions were incubated at room temperature for 1 hour. Next, 5.5 mM iodoacetamide was added and reactions were incubated on ice in the dark for 1.5 hours, dissolved in 2 volumes of 0.1 M Tris-HCl (pH 8.0) with the addition of 1 μg of trypsin (Sequencing Grade Modified, Promega) and incubated for 12 hours at 37°C. An additional 1 μg of trypsin was added and incubation at 37°C was continued for 4 more hours.

To remove urea and thiourea, tryptic peptides were bound to C18 resin (Bond Elut C18 cartridge, Agilent) in 0.1% TFA, eluted with 75% acetonitrile in 0,1% TFA, dried on a SpeedVac Vacuum Concentrator («Thermo Scientific», USA). After that, the peptides were resuspended in 20 μl 0,1% formic acid.

The proteomic analysis of peptides was performed using Ultimate 3000 RSLCnano HPLC system (Thermo Scientific, USA), connected to the Q-exact HFX mass spectrometer (Thermo Scientific, USA). 1 μl of peptide mixture was loaded onto an Acclaim µ-Precolumn enrichment column (0.5 mm x 3 mm, particle size 5 microns, Thermo Scientific, USA) at a flow rate of 10 µl/min for 5 min in isocratic mode with a 2% acetonitrile, 0.1% formic acid mobile phase.

Peptides were separated on a Peaky™ C18 HPLC column (100 microns x 30 cm, 1.9 microns particle size, Molecta, Russia) at a flow rate 0.3 μl/min using a linear (2-40%) gradient of acetonitrile in 0.1% formic acid.

MS analysis was performed on a Q-Exact HFX mass spectrometer in positive ionization mode at 2.1 kV, 240 °C using a NESI source (Thermo Scientific, USA). Panoramic scanning was performed in the mass range of 300-1500 m/z, at a resolution of 120,000. In tandem scanning, the resolution was set to 15,000 in the mass range from 100 m/z to the upper limit, which is automatically determined based on the mass of the precursor. The isolation of precursor ions was carried out within a window of ± 1 Da.

The maximum number of ions allowed for isolation in the MS2 mode was set as no more than 20, while the cut-off limit for choosing a precursor for tandem analysis was set as 400,000 units, and normalized collision energy (NCE) was 29. For tandem scanning, only ions from z = 2+ to z = 4+ were considered. The maximum accumulation time for precursor ions was 50 ms, for fragment ions 40 ms. The AGC value for precursors and fragment ions was set to 1*106 and 1*105, respectively. All the measured precursors were dynamically excluded from the tandem MS/MS analysis for 40 seconds.

For data analysis, the iBAQ algorithm was used. Comparison of relative amounts of CRISPR effector proteins and other proteins were performed using normalization on total protein iBAQ signal. For each of the 3 biological replicates, 3 technical measurements were made. For 9 normalized iBAQ signals, the average was calculated with deviations, where possible. If the signals were not detected in all 9 dimensions, then only the average is presented. The alpha subunit of RNA polymerase was used as a reference.

Proteomic assay was performed in Institute of Biomedical Chemistry, ‘Human proteome’ Core Facility Centre (Moscow, Russia)

## Data availability

The code used for the analysis is deposed on the GitHub: https://github.com/VasiliyZubarev/Thermus_CRISPR_recombination

## Accession numbers

Raw sequencing data have been deposited with the National Center for Biotechnology Information Sequence Read Archive under BioProject ID **PRJNA1004676**.

The assembly of *T. thermophilus HB27* genome was submitted to GenBank with submission ID **<u>SUB13776241</u>.**

*T. thermophilus* HB27 genome assembly GenBank: **<u>GCF_000008125.1</u>** was used for analysis of high-throughput RNA sequencing data.

## Acknowledgements

We acknowledge Skoltech Genomic Core Facility.

This work was supported by the grant of the Ministry of Science and Higher Education of the Russian Federation (075-15-2019-1661).

Y.I. was supported by FONDECYT, Chile project 1200708.

Funding for open access charge: Skolkovo Institute of Science and Technology.

## Abbreviations

CRISPR: clustered regularly interspaced short palindromic repeats
Cas: CRISPR-associated
PAM: protospacer adjacent motif
cOAs: cyclic oligoadenylates
PS: protospacer
CAR: Cas10-activating region
CARF: CRISPR-associated Rossmann Fold
LC-MS/MS: liquid chromatography coupled to tandem mass spectrometry.

