## Supplementary file 1 for "Interference Requirements of Type III CRISPR-Cas Systems from *Thermus thermophilus*"

**Supplementary figures**

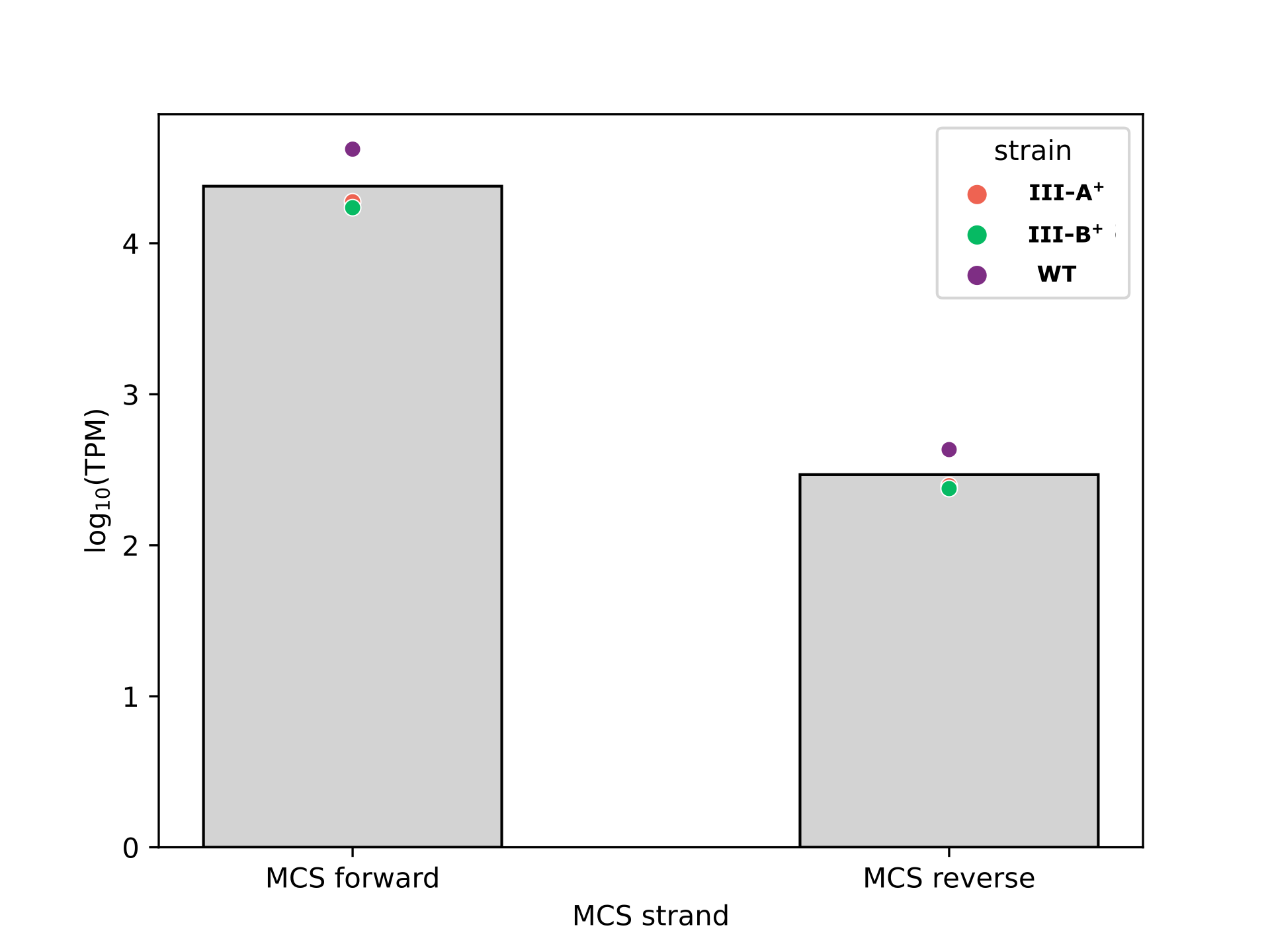

**Supplementary Figure S1. The abundance of reads corresponding to transcripts of the pMK18 polylinker site initiated from the PslpA promoter (forward) and antisense (reverse) transcripts in *T. thermophilus* HB27 wild-type, III-A^+^ and III-B^+^ strains.**
Normalized abundances of transcripts derived from both pMK18 strands mapped to a ~680 base pairs region containing the pMK18 polylinker site were determined from high throughput RNA sequencing data. The logarithms of transcript per million (TPM) values are shown as colored dots for wild-type *T. thermophilus* HB27 (purple), III-A**^+^** (red), and III-B**^+^** (green) cultures transformed with the pMK18 plasmid. Bars show means for all three strains.

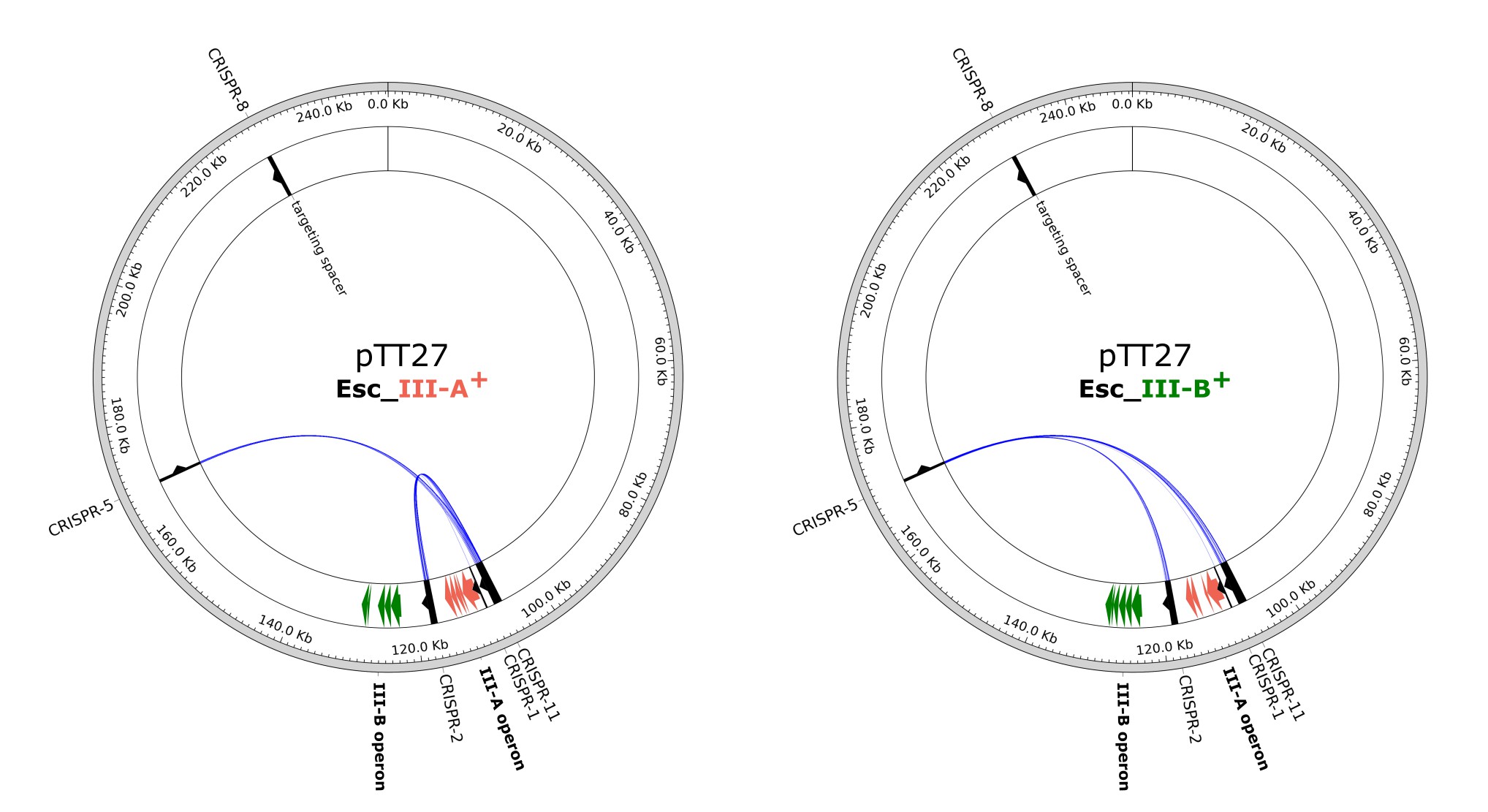

**Supplementary Figure S2. Escaper III-A^+^ and III-B^+^ clones carrying protospacer plasmids targeted by Type III immunity contain deletions of, correspondingly, III-A and III-B *cas* genes that result from recombination between flanking Type III arrays.**

Total DNA was purified from ~500 pooled kanamycin-resistant colonies obtained after transformation of III-A^+^ or III-B^+^ stains with the PS_dir plasmid. PCR amplification with primer pairs annealing outside CRISPR-5 and CRISPR-2 (Th_RecCRISPR-F2 and Th_RecCRISPR-R2 primers), CRISPR-5 and CRISPR-11 (Th_RecCRISPR-F1 and Th_RecCRISPR-R2 primers), and CRISPR-2 and CRISPR-11 (Th_RecCRISPR-F1 and Th_RecCRISPR-R1 primers, all primers are listed in Supplementary Table S1) was performed. Amplicons were pooled, subjected to Oxford Nanopore MinION sequencing and reads were mapped onto the *T. thermophilus* HB27 megaplasmid pTT27 sequence. The circos plots show genetic alterations in *T. thermophilus* observed in III-A^+^ (left) or III-B^+^ (right) transformants. The outer ring represents the pTT27 megaplasmid sequence. Type III CRISPR arrays (black, numbered according to Artamonova et al. [53]), and III-A (red) and III-B (green) *cas* operons are shown in the inner ring. Arrows indicate the direction of transcription. The spacer targeting the PS_dir protospacer is located in the CRISPR-8 array. Blue arched lines connect the boards of identified deletions.

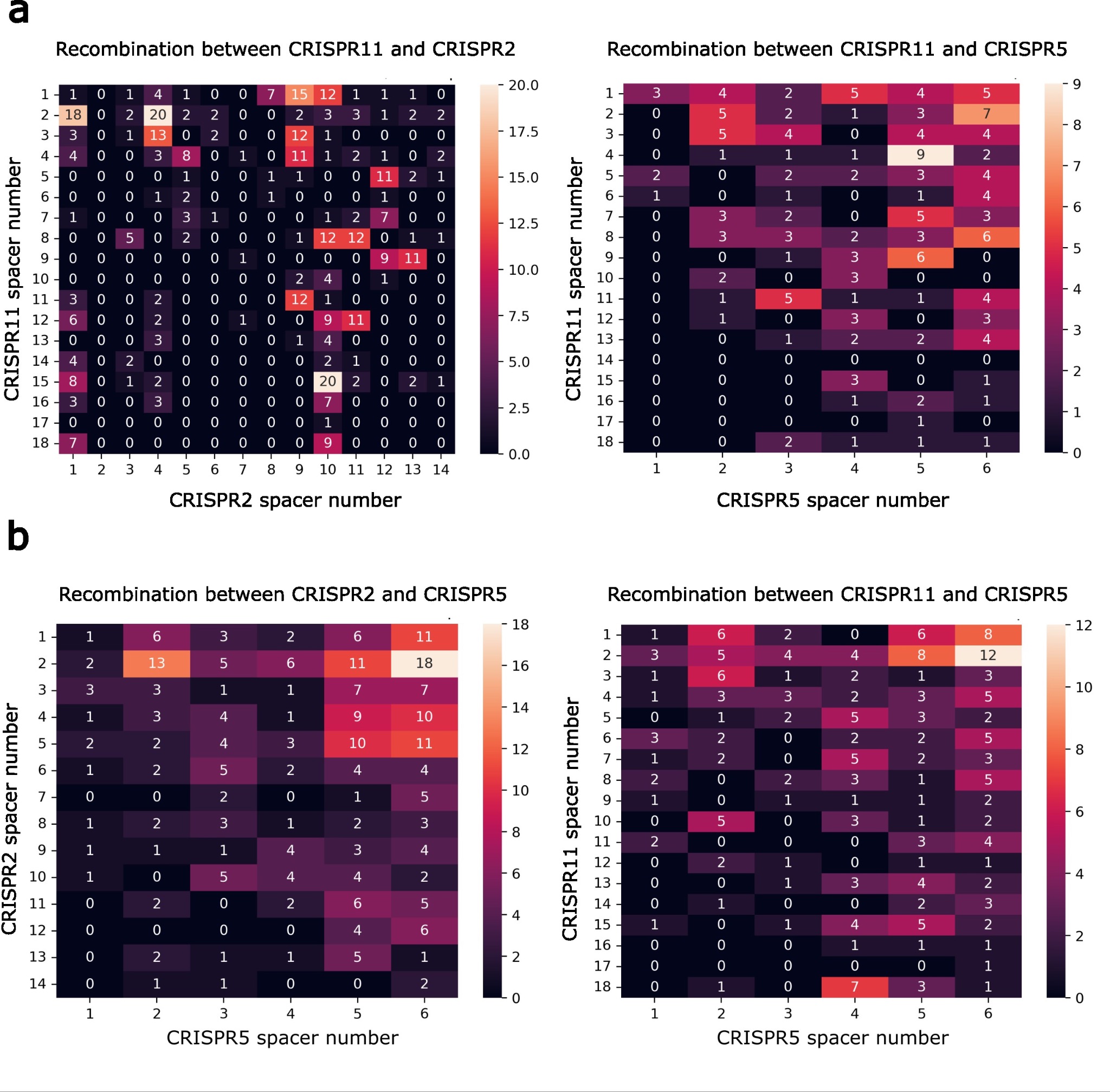

**Supplementary Figure S3. The landscape of hybrid arrays formed by escaper-generating recombination events.**

The heatmaps shows the frequency of reads corresponding to recombination events between indicated CRISPR arrays in III-A^+^ (**a**) and III-B^+^ (**b**) strains that resulted, correspondingly, in III-A and III-B *cas* operons deletions. Spacers were extracted from long sequencing reads spanning the junction points of deletions shown in Supplementary Fig. S2, and compared to the initial sets of spacers in parental CRISPR arrays using the blastn tool. Spacers in parental arrays are shown along the x- and y-axes. The spacer numbering starts from the spacer proximal to a leader sequence. Each rectangular segment on the heatmap corresponds to a possible recombination event between the arrays. Numbers and the color scheme indicate the quantity of reads corresponding to observed recombination events.

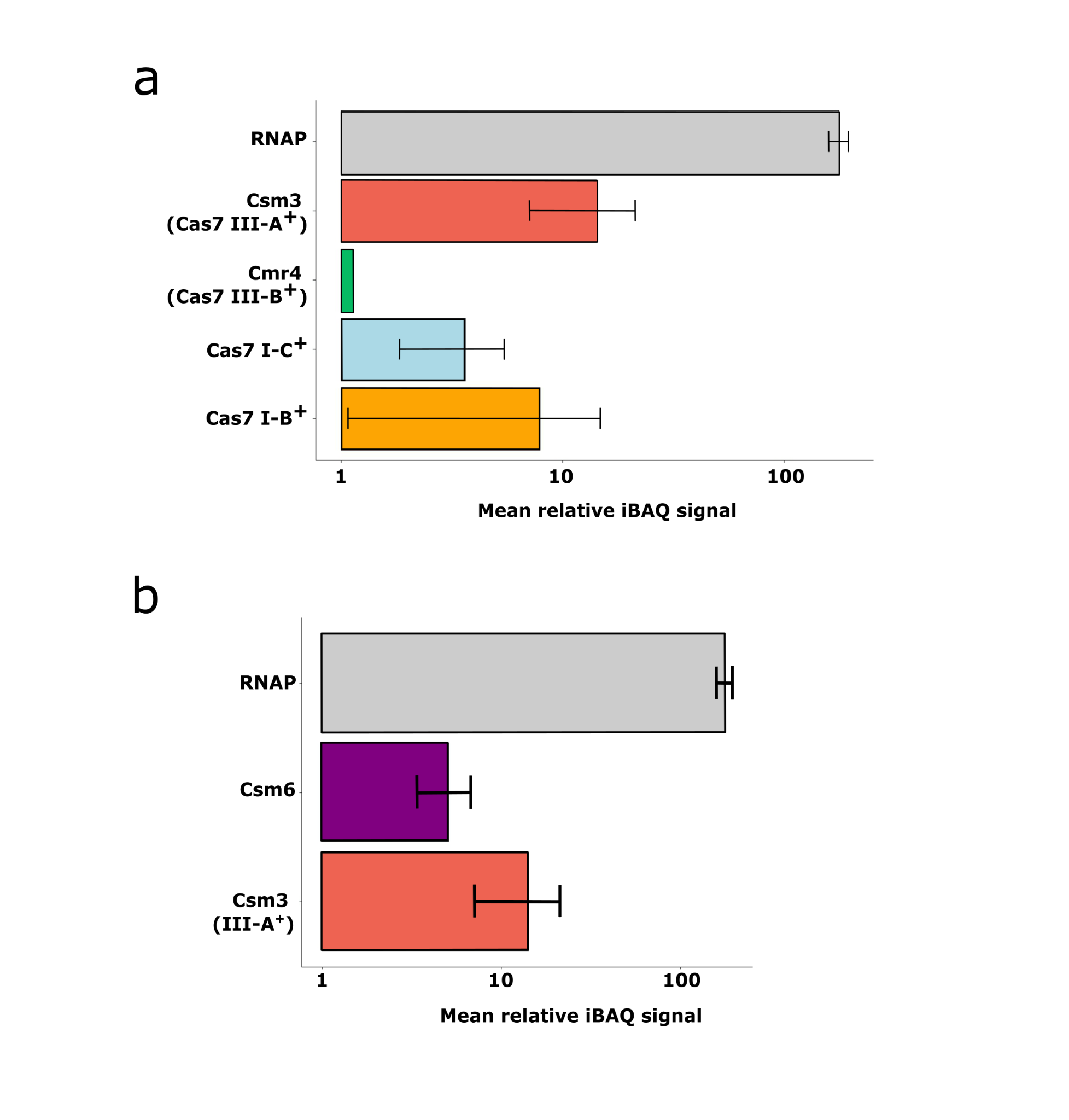

**Supplementary Figure S4. Normalized iBAQ signals for *T. thermophilus* CRISPR-Cas effector subunits (a) and CARF domain containing nuclease Csm6 (b).**
The *T. thermophilus* HB27 proteome was analyzed by liquid chromatography coupled to tandem mass spectrometry (LC-MS/MS). To evaluate the relative quantities of a particular protein in comparison to other proteins, normalization was made based on the total protein iBAQ signal. Each of the three biological replicates was subjected to three technical measurements, resulting in nine normalized iBAQ signals. Means of these signals with corresponding standart deviations are presented. In cases where signals were not detected across all nine measurements, only the calculated mean is shown. The alpha subunit of RNA polymerase was selected as a reference. In **a**, abundances of *T. thermophilus* Cas7-like effector subunits (Csm3 for Type III-A (red), Cmr4 for Type III-B (green), and homologs from Type I-C (blue) and Type I-B systems (orange)). In **b**, abundance of *T. thermophilus* Csm6 (purple), the only CARF domain containing nuclease that was detected is shown.

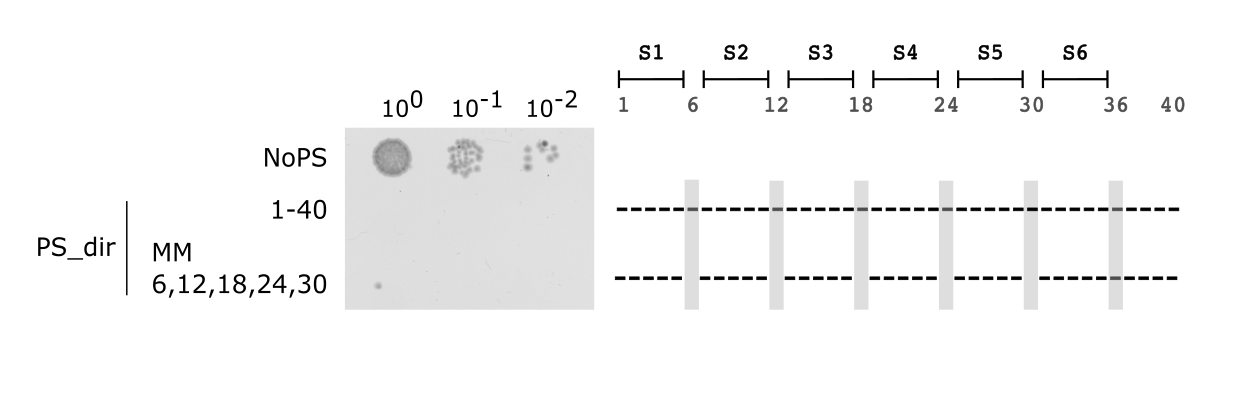

**Supplementary Figure S5. Impact of spacer-protospacer mismatches at positions +6, +12, +18, +24, and +30.**

PS_dir with five mismatches at positions +6, +12, +18, +24, and +30 was subjected to interference by the Type III-A system. Presumably, mismatches are tolerated because the corresponding positions of crRNA are flipped from the heteroduplex, interact with the Csm3 subunits of the effector and are not participating in complementary interactions with the target. See Fig. 1 legend for details.

**Supplementary tables**

**Supplementary Table S1. Oligonucleotides used in the study.**

| Name | Sequence 5′ to 3′ |
| --- | --- |
| Primers for constructions of plasmids carrying mismatched protospacers | |
| 1.8_1-4m-F | AAT TCG AGA TTC AGG ATC CAC GCA AAC TCC CTT CCT TGG GGC TTA A |
| 1.8_1-4m-R | AGC TTT AAG CCC CAA GGA AGG GAG TTT GCG TGG ATC CTG AAT CTC G |
| 1.8_1-7,23-40m-F | AAT TCG AGA AAG AGG ATC CAC GCA AAC AGG GAA GGA ACC CCG AAT A |
| 1.8_1-7,23-40m-R | AGC TTA TTC GGG GTT CCT TCC CTG TTT GCG TGG ATC CTC TTT CTC G |
| 1.8_1-8m-F | AAT TCG AGA AAG TGG ATC CAC GCA AAC TCC CTT CCT TGG GGC TTA A |
| 1.8_1-8m-R | AGC TTT AAG CCC CAA GGA AGG GAG TTT GCG TGG ATC CAC TTT CTC G |
| 1.8_1, 27-40mut_F | AAT TCG TCT TTC AGG ATC CAC GCA AAC TCC CAA GGA ACC CCG AAT A |
| 1.8_1, 27-40mut_R | AGC TTA TTC GGG GTT CCT TGG GAG TTT GCG TGG ATC CTG AAA GAC G |
| 1.8_1,2,27-40m-F | AAT TCG ACT TTC AGG ATC CAC GCA AAC TCC CAA GGA ACC CCG AAT A |
| 1.8_1,2,27-40m-R | AGC TTA TTC GGG GTT CCT TGG GAG TTT GCG TGG ATC CTG AAA GTC G |
| 1.8_17,23,29,35m-F | AAT TCC TCT TTC AGG ATC CAC CCA AAC ACC CTT GCT TGG CGC TTA A |
| 1.8_17,23,29,35m-R | AGC TTT AAG CGC CAA GCA AGG GTG TTT GGG TGG ATC CTG AAA GAG G |
| 1.8_18-23m-F | AAT TCC TCT TTC AGG ATC CAC GGT TTG ACC CTT CCT TGG GGC TTA A |
| 1.8_18-23m-R | AGC TTT AAG CCC CAA GGA AGG GTC AAA CCG TGG ATC CTG AAA GAG G |
| 1.8_19-24m-F | AAT TCC TCT TTC AGG ATC CAC GCT TTG AGC CTT CCT TGG GGC TTA A |
| 1.8_19-24m-R | AGC TTT AAG CCC CAA GGA AGG CTC AAA GCG TGG ATC CTG AAA GAG G |
| 1.8_2, 27-40mut_F | AAT TCC ACT TTC AGG ATC CAC GCA AAC TCC CAA GGA ACC CCG AAT A |
| 1.8_2, 27-40mut_R | AGC TTA TTC GGG GTT CCT TGG GAG TTT GCG TGG ATC CTG AAA GTG G |
| 1.8_22-40_new-F | AAT TCC TCT TTC AGG ATC CAC GCA AAG AGG GAA GGA ACC CCG AAT A |
| 1.8_22-40_new-R | AGC TTA TTC GGG GTT CCT TCC CTC TTT GCG TGG ATC CTG AAA GAG G |
| 1.8_22-40_rev-F | AAT TCA TTC GGG GTT CCT TCC CTC TTT GCG TGG ATC CTG AAA GAG A |
| 1.8_22-40_rev-R | AGC TTC TCT TTC AGG ATC CAC GCA AAG AGG GAA GGA ACC CCG AAT G |
| 1.8_23-40_new-F | AAT TCC TCT TTC AGG ATC CAC GCA AAC AGG GAA GGA ACC CCG AAT A |
| 1.8_23-40_new-R | AGC TTA TTC GGG GTT CCT TCC CTG TTT GCG TGG ATC CTG AAA GAG G |
| 1.8_23-40_rev_new-F | AAT TCA TTC GGG GTT CCT TCC CTG TTT GCG TGG ATC CTG AAA GAG A |
| 1.8_23-40_rev_new-R | AGC TTC TCT TTC AGG ATC CAC GCA AAC AGG GAA GGA ACC CCG AAT G |
| 1.8_24-40mut_F | AAT TCC TCT TTC AGG ATC CAC GCA AAC TGG CAA GCA ACC CGG AAT A |
| 1.8_24-40mut_R | AGC TTA TTC CGG GTT GCT TGC CAG TTT GCG TGG ATC CTG AAA GAG G |
| 1.8_26-40mut_new_F | AAT TCC TCT TTC AGG ATC CAC GCA AAC TCC GAA GGA ACC CCG AAT A |
| 1.8_26-40mut_new_R | TAT TCG GGG TTC CTT CGG AGT TTG CGT GGA TCC TGA AAG AGG AAT T |
| 1.8_26-40mut_new_R | AGC TTA TTC GGG GTT CCT TCG GAG TTT GCG TGG ATC CTG AAA GAG G |
| 1.8_27-40mut_F | AAT TCC TCT TTC AGG ATC CAC GCA AAC TCC CAA GCA ACC CGG AAT A |
| 1.8_27-40mut_F | AAT TCC TCT TTC AGG ATC CAC GCA AAC TCC CAA GGA ACC CCG AAT A |
| 1.8_27-40mut_R | AGC TTA TTC CGG GTT GCT TGG GAG TTT GCG TGG ATC CTG AAA GAG G |
| 1.8_27-40mut_R | AGC TTA TTC GGG GTT CCT TGG GAG TTT GCG TGG ATC CTG AAA GAG G |
| 1.8_5-8mut_new_F | AAT TCC TCT AAG TGG ATC CAC GCA AAC TCC CTT CCT TGG GGC TTA A |
| 1.8_5-8mut_new_R | AGC TTT AAG CCC CAA GGA AGG GAG TTT GCG TGG ATC CAC TTA GAG G |
| 1.8_5,11,17,23,29,35mut_F | AAT TCC TCT ATC AGG TTC CAC CCA AAC ACC CTT GCT TGG CGC TTA A |
| 1.8_5,11,17,23,29,35mut_R | AGC TTT AAG CGC CAA GCA AGG GTG TTT GGG TGG AAC CTG ATA GAG G |
| 1.8_7-14m-F | AAT TCC TCT TTG TCC TAG GAC GCA AAC TCC CTT CCT TGG GGC TTA A |
| 1.8_7-14m-R | AGC TTT AAG CCC CAA GGA AGG GAG TTT GCG TCC TAG GAC AAA GAG G |
| 1.8_dir_1-2-F | AAT TCG ACT TTC AGG ATC CAC GCA AAC TCC CTT CCT TGG GGC TTA A |
| 1.8_dir_1-2-R | AGC TTT AAG CCC CAA GGA AGG GAG TTT GCG TGG ATC CTG AAA GTC G |
| 1.8_dir_1-2,28-40m-F | AAT TCG ACT TTC AGG ATC CAC GCA AAC TCC CTA GGA ACC CCG AAT A |
| 1.8_dir_1-2,28-40m-R | AGC TTA TTC GGG GTT CCT AGG GAG TTT GCG TGG ATC CTG AAA GTC G |
| 1.8_dir_1-2,29-40m-R | AAT TCG ACT TTC AGG ATC CAC GCA AAC TCC CTT GGA ACC CCG AAT A |
| 1.8_dir_1-2m_F | AAT TCG ACT TTC AGG ATC CAC GCA AAC TCC CTT CCT TGG GGC TTA A |
| 1.8_dir_1-2m_R | AGC TTT AAG CCC CAA GGA AGG GAG TTT GCG TGG ATC CTG AAA GTC G |
| 1.8_dir_1-3,29-40m-F | AAT TCG AGT TTC AGG ATC CAC GCA AAC TCC CTT GGA ACC CCG AAT A |
| 1.8_dir_1-3,29-40m-R | AGC TTA TTC GGG GTT CCA AGG GAG TTT GCG TGG ATC CTG AAA CTC G |
| 1.8_dir_1-3,30-40m-F | AAT TCG AGT TTC AGG ATC CAC GCA AAC TCC CTT CGA ACC CCG AAT A |
| 1.8_dir_1-3,30-40m-R | AGC TTA TTC GGG GTT CGA AGG GAG TTT GCG TGG ATC CTG AAA CTC G |
| 1.8_dir_1-4,30-40m-F | AAT TCG AGA TTC AGG ATC CAC GCA AAC TCC CTT CGA ACC CCG AAT A |
| 1.8_dir_1-4,30-40m-R | AGC TTA TTC GGG GTT CGA AGG GAG TTT GCG TGG ATC CTG AAT CTC G |
| 1.8_dir_1-4,31-40m-F | AAT TCG AGA TTC AGG ATC CAC GCA AAC TCC CTT CCA ACC CCG AAT A |
| 1.8_dir_1-4,31-40m-R | AGC TTA TTC GGG GTT GGA AGG GAG TTT GCG TGG ATC CTG AAT CTC G |
| 1.8_dir_1-5,31-40m-F | AAT TCG AGA ATC AGG ATC CAC GCA AAC TCC CTT CCA ACC CCG AAT A |
| 1.8_dir_1-5,31-40m-R | AGC TTA TTC GGG GTT GGA AGG GAG TTT GCG TGG ATC CTG ATT CTC G |
| 1.8_dir_1-5,32-40m-F | AAT TCG AGA ATC AGG ATC CAC GCA AAC TCC CTT CCT ACC CCG AAT A |
| 1.8_dir_1-5,32-40m-R | AGC TTA TTC GGG GTA GGA AGG GAG TTT GCG TGG ATC CTG ATT CTC G |
| 1.8_dir_1-6,32-40m-F | AAT TCG AGA AAC AGG ATC CAC GCA AAC TCC CTT CCT ACC CCG AAT A |
| 1.8_dir_1-6,32-40m-R | AGC TTA TTC GGG GTA GGA AGG GAG TTT GCG TGG ATC CTG TTT CTC G |
| 1.8_dir_1-6,33-40m-F | AAT TCG AGA AAC AGG ATC CAC GCA AAC TCC CTT CCT TCC CCG AAT A |
| 1.8_dir_1-6,33-40m-R | AGC TTA TTC GGG GAA GGA AGG GAG TTT GCG TGG ATC CTG TTT CTC G |
| 1.8_dir_1-7,33-40m-F | AAT TCG AGA AAG AGG ATC CAC GCA AAC TCC CTT CCT TCC CCG AAT A |
| 1.8_dir_1-7,33-40m-R | AGC TTA TTC GGG GAA GGA AGG GAG TTT GCG TGG ATC CTC TTT CTC G |
| 1.8_dir_1-7,34-40m-F | AAT TCG AGA AAG AGG ATC CAC GCA AAC TCC CTT CCT TGC CCG AAT A |
| 1.8_dir_1-7,34-40m-R | AGC TTA TTC GGG CAA GGA AGG GAG TTT GCG TGG ATC CTC TTT CTC G |
| 1.8_dir_1-8,34-40m-F | AAT TCG AGA AAG TGG ATC CAC GCA AAC TCC CTT CCT TGC CCG AAT A |
| 1.8_dir_1-8,34-40m-R | AGC TTA TTC GGG CAA GGA AGG GAG TTT GCG TGG ATC CAC TTT CTC G |
| 1.8_dir_1-8,35-40m-F | AAT TCG AGA AAG TGG ATC CAC GCA AAC TCC CTT CCT TGG CCG AAT A |
| 1.8_dir_1-8,35-40m-R | AGC TTA TTC GGC CAA GGA AGG GAG TTT GCG TGG ATC CAC TTT CTC G |
| 1.8_dir_1m_F | AAT TCG TCT TTC AGG ATC CAC GCA AAC TCC CTT CCT TGG GGC TTA A |
| 1.8_dir_1m_R | AGC TTT AAG CCC CAA GGA AGG GAG TTT GCG TGG ATC CTG AAA GAC G |
| 1.8_dir_21-40m-F | AAT TCC TCT TTC AGG ATC CAC GCA ATG AGG GAA GGA ACC CCG AAT A |
| 1.8_dir_21-40m-R | AGC TTA TTC GGG GTT CCT TCC CTC ATT GCG TGG ATC CTG AAA GAG G |
| 1.8_dir_29-40m-F | AAT TCC TCT TTC AGG ATC CAC GCA AAC TCC CTT GGA ACC CCG AAT A |
| 1.8_dir_29-40m-R | AGC TTA TTC GGG GTT CCA AGG GAG TTT GCG TGG ATC CTG AAA GAG G |
| 1.8_dir_30-40m_F | AAT TCC TCT TTC AGG ATC CAC GCA AAC TCC CTT CGA ACC CCG AAT A |
| 1.8_dir_30-40m_R | AGC TTA TTC GGG GTT CGA AGG GAG TTT GCG TGG ATC CTG AAA GAG G |
| 1.8_dir_31-40m-F | AAT TCC TCT TTC AGG ATC CAC GCA AAC TCC CTT CCA ACC CCG AAT A |
| 1.8_dir_31-40m-R | AGC TTA TTC GGG GTT GGA AGG GAG TTT GCG TGG ATC CTG AAA GAG G |
| 1.8_dir_32-40m-F | AAT TCC TCT TTC AGG ATC CAC GCA AAC TCC CTT CCT ACC CCG AAT A |
| 1.8_dir_32-40m-R | AGC TTA TTC GGG GTA GGA AGG GAG TTT GCG TGG ATC CTG AAA GAG G |
| 1.8_dir_33-40m-F | AAT TCC TCT TTC AGG ATC CAC GCA AAC TCC CTT CCT TCC CCG AAT A |
| 1.8_dir_33-40m-R | AGC TTA TTC GGG GAA GGA AGG GAG TTT GCG TGG ATC CTG AAA GAG G |
| 1.8_dir_35-40m-F | AAT TCC TCT TTC AGG ATC CAC GCA AAC TCC CTT CCT TGG CCG AAT A |
| 1.8_dir_35-40m-R | AGC TTA TTC GGC CAA GGA AGG GAG TTT GCG TGG ATC CTG AAA GAG G |
| 1.8_rev_1-2m-F | AAT TCA TAG CCC CAA GGA AGG GAG TTT GCG TGG ATC CTG AAA GAG A |
| 1.8_rev_1-2m-F | AAT TCT AAG CCC CAA GGA AGG GAG TTT GCG TGG ATC CTG AAA GTC A |
| 1.8_rev_1-2m-R | AGC TTC TCT TTC AGG ATC CAC GCA AAC TCC CTT CCT TGG GGC TAT G |
| 1.8_rev_1-2m-R | AGC TTG ACT TTC AGG ATC CAC GCA AAC TCC CTT CCT TGG GGC TTA G |
| 1.8_rev_1-5m-F | AAT TCA TTC GCC CAA GGA AGG GAG TTT GCG TGG ATC CTG AAA GAG A |
| 1.8_rev_1-5m-F | AAT TCT AAG CCC CAA GGA AGG GAG TTT GCG TGG ATC CTG ATT CTC A |
| 1.8_rev_1-5m-R | AGC TTC TCT TTC AGG ATC CAC GCA AAC TCC CTT CCT TGG GCG AAT G |
| 1.8_rev_1-5m-R | AGC TTG AGA ATC AGG ATC CAC GCA AAC TCC CTT CCT TGG GGC TTA G |
| 1.8_rev_1-6m-F | AAT TCT AAG CCC CAA GGA AGG GAG TTT GCG TGG ATC CTG TTT CTC A |
| 1.8_rev_1-6m-R | AGC TTG AGA AAC AGG ATC CAC GCA AAC TCC CTT CCT TGG GGC TTA G |
| 1.8_rev_1m-F | AAT TCT AAG CCC CAA GGA AGG GAG TTT GCG TGG ATC CTG AAA GAC A |
| 1.8_rev_1m-R | AGC TTG TCT TTC AGG ATC CAC GCA AAC TCC CTT CCT TGG GGC TTA G |
| 1.8_rev_21-40m-F | AAT TCA TTC GGG GTT CCT TCC CTC ATT GCG TGG ATC CTG AAA GAG A |
| 1.8_rev_21-40m-R | AGC TTC TCT TTC AGG ATC CAC GCA ATG AGG GAA GGA ACC CCG AAT G |
| 1.8_rev_34-40m-F | AAT TCT AAG CCC CAA GGA AGG GAG TTT GCG TGG ATC CTC TTT CTC A |
| 1.8_rev_34-40m-F | AAT TCA TTC GGG CAA GGA AGG GAG TTT GCG TGG ATC CTG AAA GAG A |
| 1.8_rev_34-40m-R | AGC TTG AGA AAG AGG ATC CAC GCA AAC TCC CTT CCT TGG GGC TTA G |
| 1.8_rev_34-40m-R | AGC TTC TCT TTC AGG ATC CAC GCA AAC TCC CTT CCT TGC CCG AAT G |
| 1.8_rev_36-40m-F | AAT TCA TTC GCC CAA GGA AGG GAG TTT GCG TGG ATC CTG AAA GAG A |
| 1.8_rev_36-40m-R | AGC TTC TCT TTC AGG ATC CAC GCA AAC TCC CTT CCT TGG GCG AAT G |
| 1.8_rev_37-40m_F | AAT TCA TTC CCC CAA GGA AGG GAG TTT GCG TGG ATC CTG AAA GAG A |
| 1.8_rev_37-40m_R | AGC TTC TCT TTC AGG ATC CAC GCA AAC TCC CTT CCT TGG GGG AAT G |
| 1.8_rev_38-40m-F | AAT TCA TTG CCC CAA GGA AGG GAG TTT GCG TGG ATC CTG AAA GAG A |
| 1.8_rev_38-40m-R | AGC TTC TCT TTC AGG ATC CAC GCA AAC TCC CTT CCT TGG GGC AAT G |
| 1.8_rev_39-40m-F | AAT TCA TAG CCC CAA GGA AGG GAG TTT GCG TGG ATC CTG AAA GAG A |
| 1.8_rev_39-40m-R | AGC TTC TCT TTC AGG ATC CAC GCA AAC TCC CTT CCT TGG GGC TAT G |
| 1.8_rev_40m-F | AAT TCA AAG CCC CAA GGA AGG GAG TTT GCG TGG ATC CTG AAA GAG A |
| 1.8_rev_40m-R | AGC TTC TCT TTC AGG ATC CAC GCA AAC TCC CTT CCT TGG GGC TTT G |
| 1.8_rev_5,11-F | AAT TCT AAG CCC CAA GGA AGG GAG TTT GCG TGG AAC CTG ATA GAG A |
| 1.8_rev_5,11-R | AGC TTC TCT ATC AGG TTC CAC GCA AAC TCC CTT CCT TGG GGC TTA G |
| 1.8_rev_5,11,17m-F | AAT TCT AAG GCC CAA CGA AGG CAG TTT GCG TGG ATC CTG AAA GAG A |
| 1.8_rev_5,11,17m-F | AGC TTC TCT TTC AGG ATC CAC GCA AAC TGC CTT CGT TGG GCC TTA G |
| 1.8_rev_5,11,17m-F | AAT TCT AAG CCC CAA GGA AGG GAG TTT GGG TGG AAC CTG ATA GAG A |
| 1.8_rev_5,11,17m-R | AGC TTC TCT ATC AGG TTC CAC CCA AAC TCC CTT CCT TGG GGC TTA G |
| 1.8. 24-40 (F) new | AAT TCC TCT TTC AGG ATC CAC GCA AAC TGG GAA GGA ACC CCG AAT A |
| 1.8. 24-40 (R) new | AGC TTA TTC GGG GTT CCT TCC CAG TTT GCG TGG ATC CTG AAA GAG G |
| 1.8. 24-40 REV (F) new | AAT TCA TTC GGG GTT CCT TCC CAG TTT GCG TGG ATC CTG AAA GAG A |
| 1.8. 24-40 REV (R) new | AGC TTC TCT TTC AGG ATC CAC GCA AAC TGG GAA GGA ACC CCG AAT G |
| 1.8. 25-40 (F) new | AAT TCC TCT TTC AGG ATC CAC GCA AAC TCG GAA GGA ACC CCG AAT A |
| 1.8. 25-40 (R) new | AGC TTA TTC GGG GTT CCT TCC GAG TTT GCG TGG ATC CTG AAA GAG |
| 1.8. 25-40 REV (F) new | AAT TCA TTC GGG GTT CCT TCC GAG TTT GCG TGG ATC CTG AAA GAA |
| 1.8. 25-40 REV (R) new | AGC TTC TCT TTC AGG ATC CAC GCA AAC TCG GAA GGA ACC CCG AAT G |
| 1.8. 28-40 (F) new | AAT TCC TCT TTC AGG ATC CAC GCA AAC TCC CTA GGA ACC CCG AAT A |
| 1.8. 28-40 (R) new | AGC TTA TTC GGG GTT CCT AGG GAG TTT GCG TGG ATC CTG AAA GAG G |
| 1.8. 28-40 REV (F) new | AAT TCA TTC GGG GTT CCT AGG GAG TTT GCG TGG ATC CTG AAA GAG A |
| 1.8. 28-40 REV (R) new | AGC TTC TCT TTC AGG ATC CAC GCA AAC TCC CTA GGA ACC CCG AAT G |
| 1.8. 29-40 (F) new | AAT TCC TCT TTC AGG ATC CAC GCA AAC TCC CTA CGA ACC CCG AAT A |
| 1.8. 29-40 (R) new | AGC TTA TTC GGG GTT CGT AGG GAG TTT GCG TGG ATC CTG AAA GAG G |
| 1.8. 29-40 REV (F) new | AAT TCA TTC GGG GTT CGT AGG GAG TTT GCG TGG ATC CTG AAA GAG A |
| 1.8. 29-40 REV (R) new | AGC TTC TCT TTC AGG ATC CAC GCA AAC TCC CTA CGA ACC CCG AAT G |
| 1.8rev_1-3mut-F | AAT TCT AAG CCC CAA GGA AGG GAG TTT GCG TGG ATC CTG AAA CTC A |
| 1.8rev_1-3mut-R | AGC TTG AGT TTC AGG ATC CAC GCA AAC TCC CTT CCT TGG GGC TTA G |
| 1.8rev_1-4mut-F | AAT TCT AAG CCC CAA GGA AGG GAG TTT GCG TGG ATC CTG AAT CTC A |
| 1.8rev_1-4mut-R | AGC TTG AGA TTC AGG ATC CAC GCA AAC TCC CTT CCT TGG GGC TTA G |
| 1.8rev_1-7mut-F | AAT TCT AAG CCC CAA GGA AGG GAG TTT GCG TGG ATC CTC TTT CTC A |
| 1.8rev_1-7mut-R | AGC TTG AGA AAG AGG ATC CAC GCA AAC TCC CTT CCT TGG GGC TTA G |
| 1.8rev_1,28-40m_F | AAT TCA TTC GGG GTT CCT AGG GAG TTT GCG TGG ATC CTG AAA GAC A |
| 1.8rev_1,28-40m_R | AGC TTG TCT TTC AGG ATC CAC GCA AAC TCC CTA GGA ACC CCG AAT G |
| 1.8rev_1,29-40m_F | AAT TCA TTC GGG GTT CCA AGG GAG TTT GCG TGG ATC CTG AAA GAC A |
| 1.8rev_1,29-40m_R | AGC TTG TCT TTC AGG ATC CAC GCA AAC TCC CTT GGA ACC CCG AAT G |
| 1.8rev_23-40mut_F | AAT TCA TTC GGG GTT CCT TCC CTG TTT GCG TGG ATC CTG AAA GAG A |
| 1.8rev_23-40mut_R | AGC TTC TCT TTC AGG ATC CAC GCA AAC AGG GAA GGA ACC CCG AAT G |
| 1.8rev_25-40mut_F | AAT TCA TTC GGG GTT CCT TCC GAG TTT GCG TGG ATC CTG AAA GAG A |
| 1.8rev_26-40mut_F | AAT TCA TTC GGG GTT CCT TCG GAG TTT GCG TGG ATC CTG AAA GAG A |
| 1.8rev_26-40mut_R | AGC TTC TCT TTC AGG ATC CAC GCA AAC TCC GAA GGA ACC CCG AAT G |
| 1.8rev_27-40mut_F | AAT TCA TTC GGG GTT CCT TGG GAG TTT GCG TGG ATC CTG AAA GAG A |
| 1.8rev_27-40mut_R | AGC TTC TCT TTC AGG ATC CAC GCA AAC TCC CAA GGA ACC CCG AAT G |
| 1.8rev_29-40mut_F | AAT TCA TTC GGG GTT CCA AGG GAG TTT GCG TGG ATC CTG AAA GAG A |
| 1.8rev_29-40mut_R | AGC TTC TCT TTC AGG ATC CAC GCA AAC TCC CTT GGA ACC CCG AAT G |
| 1.8rev_30-40mut_F | AAT TCA TTC GGG GTT CGA AGG GAG TTT GCG TGG ATC CTG AAA GAG A |
| 1.8rev_30-40mut_R | AGC TTC TCT TTC AGG ATC CAC GCA AAC TCC CTT CGA ACC CCG AAT G |
| 1.8rev_31-40mur_F | AAT TCA TTC GGG GTT GGA AGG GAG TTT GCG TGG ATC CTG AAA GAG A |
| 1.8rev_31-40mut_R | AGC TTC TCT TTC AGG ATC CAC GCA AAC TCC CTT CCA ACC CCG AAT G |
| 1.8rev_32-40mut_F | AAT TCA TTC GGG GTA GGA AGG GAG TTT GCG TGG ATC CTG AAA GAG A |
| 1.8rev_32-40mut_R | AGC TTC TCT TTC AGG ATC CAC GCA AAC TCC CTT CCT ACC CCG AAT G |
| 1.8rev_33-40mut_F | AAT TCA TTC GGG GAA GGA AGG GAG TTT GCG TGG ATC CTG AAA GAG A |
| 1.8rev_33-40mut_R | AGC TTC TCT TTC AGG ATC CAC GCA AAC TCC CTT CCT TCC CCG AAT G |
| 1.8rev_34-40mut_F | AAT TCA TTC GGG CAA GGA AGG GAG TTT GCG TGG ATC CTG AAA GAG A |
| 1.8rev_34-40mut_R | AGC TTC TCT TTC AGG ATC CAC GCA AAC TCC CTT CCT TGC CCG AAT G |
| 1.8rev_35-40mut_F | AAT TCA TTC GGC CAA GGA AGG GAG TTT GCG TGG ATC CTG AAA GAG A |
| 1.8rev_35-40mut_R | AGC TTC TCT TTC AGG ATC CAC GCA AAC TCC CTT CCT TGG CCG AAT G |
| 1.8rev_5,11,17,23mut-F | AAT TCT AAG CCC CAA GGA AGG GTG TTT GGG TGG AAC CTG ATA GAG A |
| 1.8rev_5,11,17,23mut-R | AGC TTC TCT ATC AGG TTC CAC CCA AAC ACC CTT CCT TGG GGC TTA G |
| 1.8rev_5,17mut-F | AAT TCT AAG CCC CAA GGA AGG GAG TTT GGG TGG ATC CTG ATA GAG A |
| 1.8rev_5,17mut-R | AGC TTC TCT ATC AGG ATC CAC CCA AAC TCC CTT CCT TGG GGC TTA G |
| 5’Handle+1.8dir_F | AAT TCA TTG CGA CCT CTT TCA GGA TCC ACG CAA ACT CCC TTC CTT GGG GCT TAA |
| 5’Handle+1.8dir_R | AGC TTT AAG CCC CAA GGA AGG GAG TTT GCG TGG ATC CTG AAA GAG GTC GCA ATG |
| 5’Handle+1.8dir1-7m_F | AAT TCA TTG CGA CGA GAA AGA GGA TCC ACG CAA ACT CCC TTC CTT GGG GCT TAA |
| 5’Handle+1.8dir1-7m_R | AGC TTT AAG CCC CAA GGA AGG GAG TTT GCG TGG ATC CTC TTT CTC GTC GCA ATG |
| 5’Handle+1.8dir1-8m_F | AAT TCA TTG CGA CGA GAA AGT GGA TCC ACG CAA ACT CCC TTC CTT GGG GCT TAA |
| 5’Handle+1.8dir1-8m_R | AGC TTT AAG CCC CAA GGA AGG GAG TTT GCG TGG ATC CAC TTT CTC GTC GCA ATG |
| dir 1,2,10,16,22,28,34-F | AAT TCG ACT TTC AGC ATC CAG GCA AAG TCC CTA CCT TGC GGC TTA A |
| dir 1,2,10,16,22,28,34-R | AGC TTT AAG CCG CAA GGT AGG GAC TTT GCC TGG ATG CTG AAA GTC G |
| DIR 1,2,5,11,17,23,29-F | AAT TCG ACT ATC AGG TTC CAC CCA AAC ACC CTT GCT TGG GGC TTA A |
| DIR 1,2,5,11,17,23,29-R | AGC TTT AAG CCC CAA GCA AGG GTG TTT GGG TGG AAC CTG ATA GTC G |
| dir 1,2,6,12,18,24,30-F | AAT TCG ACT TAC AGG AAC CAC GGA AAC TGC CTT CGT TGG GGC TTA A |
| dir 1,2,6,12,18,24,30-R | AGC TTT AAG CCC CAA CGA AGG CAG TTT CCG TGG TTC CTG TAA GTC G |
| dir 1,2,8,14,20,26,32 -R | AGC TTT AAG CCC CTA GGA ACG GAG TAT GCG TCG ATC CAG AAA GTC G |
| dir 1,2,8,14,20,26,32-F | AAT TCG ACT TTC TGG ATC GAC GCA TAC TCC GTT CCT AGG GGC TTA A |
| DIR 3,9,15,21,27-F | AAT TCC TGT TTC ACG ATC CTC GCA ATC TCC CAT CCT TGG GGC TTA A |
| DIR 3,9,15,21,27-R | AGC TTT AAG CCC CAA GGA TGG GAG ATT GCG AGG ATC GTG AAA CAG G |
| dir 6,12,18,24,30-F | AAT TCC TCT TAC AGG AAC CAC GGA AAC TGC CTT CGT TGG GGC TTA A |
| dir 6,12,18,24,30-R | AGC TTT AAG CCC CAA CGA AGG CAG TTT CCG TGG TTC CTG TAA GAG G |
| dir 8,14,20,26,32-F | AAT TCC TCT TTC TGG ATC GAC GCA TAC TCC GTT CCT AGG GGC TTA A |
| dir 8,14,20,26,32-R | AGC TTT AAG CCC CTA GGA ACG GAG TAT GCG TCG ATC CAG AAA GAG G |
| Primers for analysis of escaper diversity | |
| Th_RecCRISPR-F1 | ATT GAG GTG CAG GCT CA |
| Th_RecCRISPR-F2 | GGT AGG CCT CTT CCA TTT G |
| Th_RecCRISPR-R1 | GCC GGG AAG GAG TAG AAG A |
| Th_RecCRISPR-R2 | GGG GTT CTT GTC CCT CC |
| Primers for construction of III-A^+^ strain with mutated HD domain of Cas10 | |
| Cas10_HDcheck_F | GAG GTC CAG GTG CTG TG |
| Cas10_HDcheck_R | TTG AGG GCC CAC CTA C |
| Left_flank_fw | GGT CGA CTC TAG AGG ATC TAC TAG TCA TAT GGA TAT GGG AAA GCG TCT CTA TGC CGT |
| Left_flank_rv | TGG CCG AGG AGC AGG ACT AAT CAC AGC ACC TGG ACC TCC C |
| Right_flank_fw | CTT AAG GTT TCT GTT ATA CTC CCG GGG GGC CCC GCG TGC ACA CTT CA |
| Right_flank_rv | TCG GTA CCC GGG GAT CCG ATA GGA GGA GGT AGA ACT TGC C |
| Mut_fw | GCG GGA CTT TTG GCG GCG GTG GGC AAG CTC TAT TCC CGC G |
| Mut_rv | CTT GCC CAC CGC CGC CAA AAG TCC CGC CAG GGC CAC GCT C |
| Primers for construction of *Δcsx1, Δcsm6, Δcsx1Δcsm6* mutants | |
| P0152_LF_EcoRI_fwd | GCC TGG GAA TTC CAC GTC CTG GTG AGG GAG |
| P0152_LF_rev | GCT TCA CCT CCT AGA GGG GC |
| P0152_RF_BamHI_fwd | GGA GGC TAG AGC CCC TCT AGG AGG TGA AGC GGA TCC CGC CCT GGT CCC GGC CTT G |
| P0152_RF_rev | GGG GGT TTG TGA GAA AGA TG |
| p0142_LF_fwd | CAT ATG GAT ATC GGA TCC CCA CGT CTC GGG CCG GGT CCG GGC G |
| P0142_LF_NotI_rev | CTA GAA CCT GGG CGG CCG CCC ATT ATC CCG GGC CTC AGC AAA G |
| P0142_RF_NotI_fwd | CGG GAT AAT GGG CGG CCG CCC AGG TTC TAG GCC ATG ACC CTG |
| p0142_RF_rev | TGA ATT CGA GCT CGG TAC CCC CCG GCG AGC CGC ACC GCC ACC A |
| hygR_fwd | CTA TTC CTT TGC CCT CGG ACG AG |
| hygR_rev | CCC GGG GGG AGT ATA ACA GAA AC |
| dP0152_fwd | GTG GAC GGC TAC CCC TTA G |
| dP0152_rev | CAA ATG GAA GAG GCC TAC CGC |
| dP0142_fwd | TCG CCT TTT TGG ACC AAG CG |
| dP0142_rev | TCC AGG GAG TAA AAG AAC ACC G |
| Primers for determination of plasmid and chromosomal DNA concentration | |
| kanF(qPCR) | GCA AGG ACC GAC AAC ATT TC |
| kanR(qPCR) | CGA AGC GCT CGT CGT ATA A |
| rho_Thermus_HB27_F | CTT GTT GTT CTT GGT GCG GG |
| rho_Thermus_HB27_R | TTG ACA TCC TCA AGT CCG GC |

**Supplementary Table S2. Mutations detected in randomly chosen kanamycin-resistant clones formed after transformation of protospacer-bearing plasmids in III-A^+^ or III-B^+^ strains.**

| Escaper # | Altered gene | Alteration type |  |
| --- | --- | --- | --- |
| *Δcmr4*_1 | *cas10*/*csm1* | point mutation, frameshift | del G (Val393) |
| *Δcmr4*_2 | *cas10*/*csm1* | point mutation, frameshift | del G (Val393) |
| *Δcmr4_3* | *csm* operon | deletion of 11322 nucleotides |  |
| *Δcmr4_4* | *csm* operon | deletion of 10796 nucleotides |  |
| *Δcmr4_5* | *csm* operon | deletion of 89984 nucleotides |  |
| *Δcmr4_6* | *csm* operon | deletion of 10646 nucleotides |  |
| *Δcmr4_7* | *csm* operon | deletion of 89984 nucleotides |  |
| *Δcmr4_8* | *csm* operon | deletion of 11322 nucleotides |  |
| *Δcmr4_9* | *cas10*/*csm1* | point mutation, frameshift | del G (Val393) |
| *Δcmr4_10* | *cas10*/*csm1* | point mutation, frameshift | del C (Leu306) |
| *Δcsm3*_1 | *cas10*/*csm1* | point mutation, frameshift | del G (Gly254) |
| *Δcsm3*_2 | *cas10*/*csm1* | point mutation, frameshift | del C (Leu480) |
| *Δcsm3*_3 | *cas10*/*csm1* | point mutation, frameshift | del G (Asp381) |
| *Δcsm3*_5 | *cmr4* | point mutation, missense mutation | T>C (Leu9Pro) |
| *Δcsm3*_6 | *cmr* operon | deletion of 52276 nucleotides |  |
| *Δcsm3*_7 | *cas10*/*cmr2* | point mutation, frameshift | del C (Leu480) |
| *Δcsm3*_8 | *cas10*/*cmr2* | point mutation, frameshift | dup G (Asp381) |
| *Δcsm3*_11 | *cas10*/*cmr2* | point mutation, frameshift | del C (Leu480) |
| *Δcsm3*_12 | *cmr* operon | deletion of 52125 nucleotides |  |
| *Δcsm3*_13 | *cas10*/*cmr2* | point mutation, frameshift | dup G (Asp381) |
